# Interrupting ELMSAN1 repression of nuclear acetyl-CoA production therapeutically reprograms cancer cells

**DOI:** 10.1101/2025.04.25.650293

**Authors:** Ting Zhao, Lingli He, Lai Ping Wong, Shenglin Mei, Jun Xia, Yanxin Xu, Jonathan G. Van Vranken, Michael Mazzola, Lei Chen, Catherine Rhee, Tiancheng Fang, Tsuyoshi Fukushima, Leanne C. Sayles, Matthew Diaz, J. Alex B. Gibbons, Raul Mostoslavsky, Steven P. Gygi, Zhixun Dou, David B. Sykes, Ruslan I. Sadreyev, E. Alejandro Sweet-Cordero, David T. Scadden

**Affiliations:** Center for Regenerative Medicine, Massachusetts General Hospital, Boston, MA, USA; Harvard Stem Cell Institute, Cambridge, MA, USA; Department of Stem Cell and Regenerative Biology, Harvard University, Cambridge, MA, USA; Department of Molecular Biology, Massachusetts General Hospital, Harvard Medical School, Boston, MA, USA; Department of Cell Biology, Harvard Medical School, Boston, MA, USA; Division of Hematology and Oncology, Department of Pediatrics, University of California San Francisco, San Francisco, CA, USA; Krantz Family Center for Cancer Research, Massachusetts General Hospital, Harvard Medical School, Boston, MA, USA; Department of Pathology, Massachusetts General Hospital, Harvard Medical School, Boston, MA, USA

## Abstract

Metabolites are essential substrates for epigenetic modifications. Although nuclear acetyl-CoA constitutes a small fraction of the whole cell pool, it regulates cell fate by locally providing histone acetylation substrate. Here, we combined phenotypic chemical screen and genome-wide CRISPR screen to demonstrate a nucleus-specific acetyl-CoA regulatory mechanism that can be modulated to achieve therapeutic cancer cell reprogramming. While previously thought that nucleus-localized pyruvate dehydrogenase complex (nPDC) is constitutively active, we found that nPDC is constitutively inhibited by the nuclear protein ELMSAN1 through direct interaction. Pharmacologic inhibition of the ELMSAN1-nPDC interaction derepressed nPDC activity, enhancing nuclear acetyl-CoA generation and reprogramming cancer cells to a postmitotic state with diminished cell-of-origin signatures. Reprogramming was synergistically enhanced by histone deacetylase 1/2 inhibition, resulting in inhibited tumor growth, durably suppressed tumor-initiating ability, and improved survival in multiple cancer types *in vivo*, including therapy-resistant sarcoma patient-derived xenografts and carcinoma cell line xenografts. Our findings highlight the potential of targeting ELMSAN1-nPDC as epigenetic cancer therapy.

## INTRODUCTION

Hallmarks of cancer include deregulating cellular metabolism and non-mutational epigenetic reprogramming.^1–4^ Mutations altering epigenetic regulation are prominent cancer drivers (e.g., *DNMT3A*, *EZH2*, and *TET2*) and metabolite participation in cancer epigenetics is evident in *IDH1/2* mutant malignancies.^5–10^ During cancer progression or therapies, cancer cells undergo non-genetic state changes that usually lead to worsening outcomes, such as epithelial-mesenchymal transition (EMT) or “transdifferentiation” to a drug-resistant state.^3,11–13^ Therefore, targeting mutated metabolic/epigenetic modifiers or dysregulated cell states is a promising strategy for cancer therapy. The most notable example is differentiation therapy, which coaxes cells to resume the differentiation process to a terminal mature stage and is now being applied to cure a subset of myeloid leukemias.^14,15^ Cell state can also be modulated by targeting non-mutated epigenetic modifiers as with histone deacetylase inhibitors (HDACi).^16^ However, epigenetic targeting of cancer cell states faces challenges, including selectivity to limited cancer types, generally low efficacy on solid tumors, and unsustained effects upon drug removal.^17,18^

We sought to test whether an unbiased phenotypic small molecule screen could reveal new metabolic/epigenetic targets with perhaps fewer liabilities. We screened using osteosarcoma, a mesenchymal malignancy and the most common primary cancer in bone, with a high prevalence in adolescents and an unchanged survival rate for over four decades.^19–21^ Targeted therapy and immunotherapy have had limited effects on osteosarcoma,^22^ largely due to highly unstable and complex genetics, including *TP53* mutations or structural rearrangements in 80-90% of patients.^19,23^ Osteosarcoma exemplifies a cancer type where identifying genotype-targeted treatments is highly difficult. Successful identification of phenotype-targeted drugs and targets in osteosarcoma would allow us to potentially translate to other cancer types.

In this study, we identified compounds in phenotypic small molecule screens and studied the mechanisms of action by combining genome-wide CRISPR screens and biochemical assays. We found that a nucleus-localized component of the mitotic deacetylase complex (MiDAC), ELMSAN1, had the additional function of constitutively repressing nPDC. Pharmacologic disruption of the ELMSAN1-PDC interaction derepressed nPDC, increased nuclear acetyl-CoA, and reprogrammed the cancer cell state. This process synergized with HDAC1/2 inhibition. The result is a targeted strategy to therapeutically reprogram multiple cancer types, including patient-derived xenograft (PDX) models of intractable sarcomas.

## RESULTS

### Therapeutic cancer cell reprogramming by small molecules

Osteosarcoma exhibits a stem/progenitor-like phenotype with high proliferation and SP7 expression that can be recapitulated by a genetically engineered mouse model (GEMM) with *Tp53* and *Rb* conditional knockout in *Sp7*-expressing osteoprogenitors^24–26^ (Figure S1A). Osteoprogenitor cells normally develop into non-proliferative osteoblasts capable of calcium deposition detected by alizarin red S (ARS) staining. Accordingly, we developed a high-throughput phenotypic screening platform in GEMM cells combining optimized ARS and viability quantification to assess the loss of stemness and proliferation phenotypes at two concentrations (Figures 1A and S1B; STAR Methods). Among 3,046 compounds, we found that histone deacetylase inhibitors (HDACi), such as MGCD0103, increased ARS and decreased viability simultaneously (Figure S1C; Table S1). But when translating to human osteosarcoma patient-derived xenograft cells (PDXC), MGCD0103 exhibited a much weaker effect on ARS. We then conducted a second ARS screen with MGCD0103 in combination to assess the top 76 primary screening candidates at a 9-dose gradient in human osteosarcoma cells (Figure 1A). We generated the dose-response curve of all compounds and found that ISX9 induced the highest ARS at relatively low concentrations (Figure 1B). ISX9 also scored highest in a qPCR screen to assess cell state change among primary screening candidates and curated cell fate-changing compounds^27^ (Figure S1D). We used multiple PDXC lines to validate that ISX9 alone robustly induced ARS and reduced cell growth independent of HDACi (Figure 1C). The target of ISX9 is not well defined, but it is reported to promote neurogenic differentiation in mouse neural stem cells.^28^

**Figure 1.**
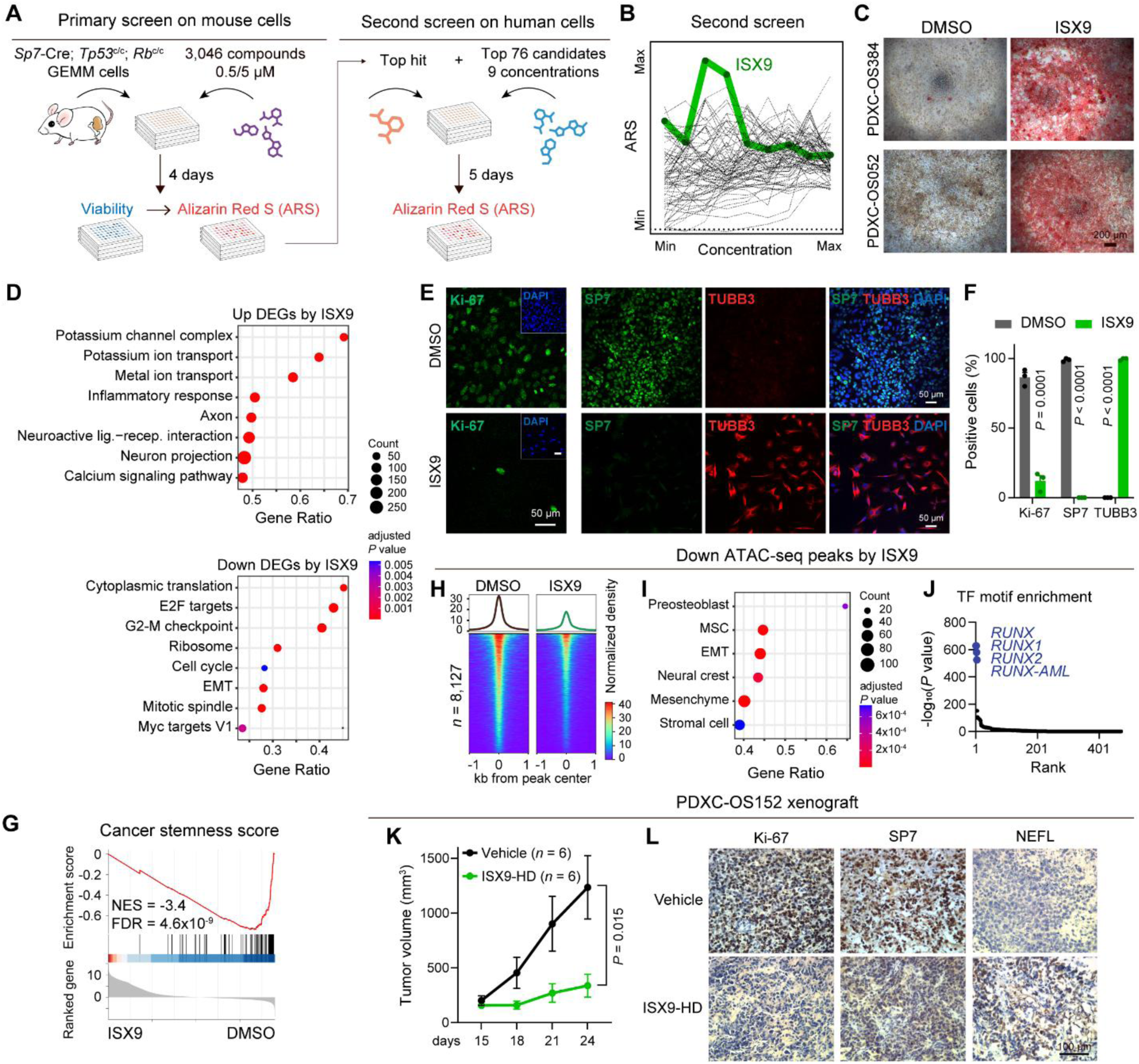
Therapeutic cancer cell reprogramming by small molecules. (A) Schematic of phenotypic small molecule screen. (B) Dose-response curves in the second ARS screen on the top of MGCD0103 in MG63 cells. A low ARS signal was observed under high doses of ISX9 because of cell death. (C) Representative pictures of ARS staining under ISX9 treatment for 8 days in two PDXC lines. Scale bar, 200 µm. (D) Enrichment analysis of DEGs by 100 µM ISX9 treatment for 6 days. (E) Immunofluorescence images of Ki-67 and SP7+TUBB3 co-staining under DMSO and 100 µM ISX9 treatment. Scale bar, 50 µm. (F) Quantification of Ki-67, SP7, and TUBB3 positive cells. *n* = 3. (G) Gene set enrichment analysis (GSEA) of cancer stemness score^29^ by 100 µM ISX9 treatment. NES: normalized enrichment score. FDR: false discovery rates. (H-J) Heatmaps and average profiles (H), enrichment analysis (I), and TF binding motif enrichment ranked by *P*-values (J) of decreased ATAC-seq peaks by 100 µM ISX9. *n* = 2. (K, L) Tumor growth (*n* = 6, K) and immunochemistry staining of Ki-67, SP7, and NEFL in tumors (L)of PDXC-OS152 xenograft under vehicle and ISX9-HD (50 mg/kg, twice daily) treatment. Scale bar, 100 µm. Data are presented as mean ± SEM. Experiments were done in PDXC-OS052 unless otherwise indicated. Statistical significance was determined using Benjamini-Hochberg method (D, I), binomial test (J), or two-tailed unpaired t-test (F, K). See also Figure S1 and Tables S1-2.

To better understand the ISX9-induced phenotype, we performed RNA-seq and ATAC-seq in osteosarcoma PDXCs after six days of treatment (Table S2). Differentially expressed gene (DEG) analysis showed that the most dramatic change was the downregulation of genes enriched in E2F targets and G2-M checkpoint (Figure 1D), such as *MKI67*, *AURKA*, *FOXM1*, and *CCNA2* (Figure S1E). This result was validated by a significant reduction of Ki-67 staining (Figures 1E and 1F). Reduced cell cycling was associated with upregulated *CDKN1A* (*P21*) (Figure S1E) but not cellular senescence (Figure S1F). ISX9 also significantly suppressed the cancer stemness score^29^ (Figure 1G), supporting the loss of the stemness phenotype. Notably, ISX9 downregulated genes enriched in EMT (Figure 1D), such as transcription factor (TF) *TWIST1*, *ZEB2*, and *SNAI2* and osteogenic genes, including *SP7*, *BGLAP*, and *COL1A1* (Figure S1E). Immunofluorescence confirmed the complete loss of SP7 protein upon ISX9 treatment (Figures 1E and 1F). Correspondingly, ATAC-seq showed that the decreased accessible regions were highly enriched for EMT and mesenchymal tissue signatures (Figures 1H and 1I; Table S2). The binding motifs of the RUNX family, including the master osteogenic TF *RUNX2*, were mostly and uniquely enriched in reduced accessible regions by ISX9 (Figures 1J and S1G), supporting an epigenetic alteration of the cell-of-origin signature.

Strikingly, ISX9 upregulated genes enriched in ion transport, neuron projection, and axon (Figure 1D), such as *NEUROD1*, *POU3F2*, *TUBB3*, *NEFL*, and *NEFM* (Figure S1E). Immunofluorescence showed that ∼100% of ISX9-treated cells were TUBB3^+^ with 100% SP7^−^ and ∼90% Ki-67^−^ (Figures 1E and 1F). The highly efficient and rapid cell state transition argued against the outgrowth of a preexisting subclone that is often seen in the setting of drug-induced resistance.^30^ ATAC-seq also showed open chromatin accessibility in neuronal genes upon ISX9 treatment (Figures S1H and S1I). However, the levels of neuronal gene activation were low, and neuronal TF protein expression was minimal by immunofluorescence.

In a xenograft model of osteosarcoma PDXC *in vivo*, ISX9 was administered at a high dose (ISX9-HD, 50 mg/kg, twice daily), which differed from the low-dose ISX9 (ISX9-LD, 25-40 mg/kg, once daily) used in combination with MGCD0103. ISX9-HD significantly reduced tumor growth and prolonged mouse survival without affecting body weight (Figures 1K and S1J-S1L). Immunohistochemistry showed a similar phenotypic change *in vivo*, including the downregulation of Ki-67, SP7, and upregulation of NEFL in tumors from ISX9-HD treated mice (Figure 1L). In summary, we employed phenotypic screens to identify small molecule ISX9 that reprogrammed a mesenchymal-neural-like transition (MNLT) in osteosarcoma cells to exhibit several clinically therapeutic phenotypes, including postmitotic state, reduced stemness, diminished mesenchymal and cell-of-origin identity, and delayed tumor growth. A subtle “neural-like” phenotype is observed, likely being a “passenger” rather than a “driver” event and attributed to the ARS screen since the critical enzyme in calcium deposition, alkaline phosphatase, also plays a role in promoting axonal growth in neurons.^31,32^

### Genome-wide CRISPR activation screen identifies ELMSAN1 as the reprogramming mediator

Next, we used ISX9 as a tool compound to identify its mechanism of action (MOA) in inducing the therapeutic phenotype. ISX9 was reported to activate *Neurod1* through Ca^2+^-CaMKII-mediated nuclear export of HDAC5 in neural progenitor cells,^28^ but we did not observe this mechanism in osteosarcoma (Figure S2A). Attempts to identify ISX9 target proteins using affinity chromatography^33^ and proteome integral stability alteration (PISA)^34^ were unsuccessful despite further confirmation of MGCD0103 targets (Figure S2B).

We then conducted a genome-wide CRISPR activation (CRISPRa) screen to identify genes whose overexpression blunted the ISX9 effect in osteosarcoma 143B cells (Figures 2A, S2C, and S2D; Table S3). Strikingly, *ELMSAN1* (also named *MIDEAS*) ranked highest in antagonizing ISX9 (Figures 2B and S2E). We validated that *ELMSAN1* overexpression (OE) increased, while knockdown (KD) decreased the half-maximal inhibitory concentration (IC_50_) of ISX9 (Figures 2C and S2F). *ELMSAN1* similarly affected the sensitivity of HDACi, including MGCD0103 and other clinically tested HDACi (Figures S2G and S2H). Additionally, DepMap analysis showed a significant correlation between *ELMSAN1* expression and sensitivity to ISX9 in various cancer types, which were consistently negative (Figure 2D). These data imply a convergent role of ELMSAN1 on the effect of ISX9 and HDACi.

**Figure 2.**
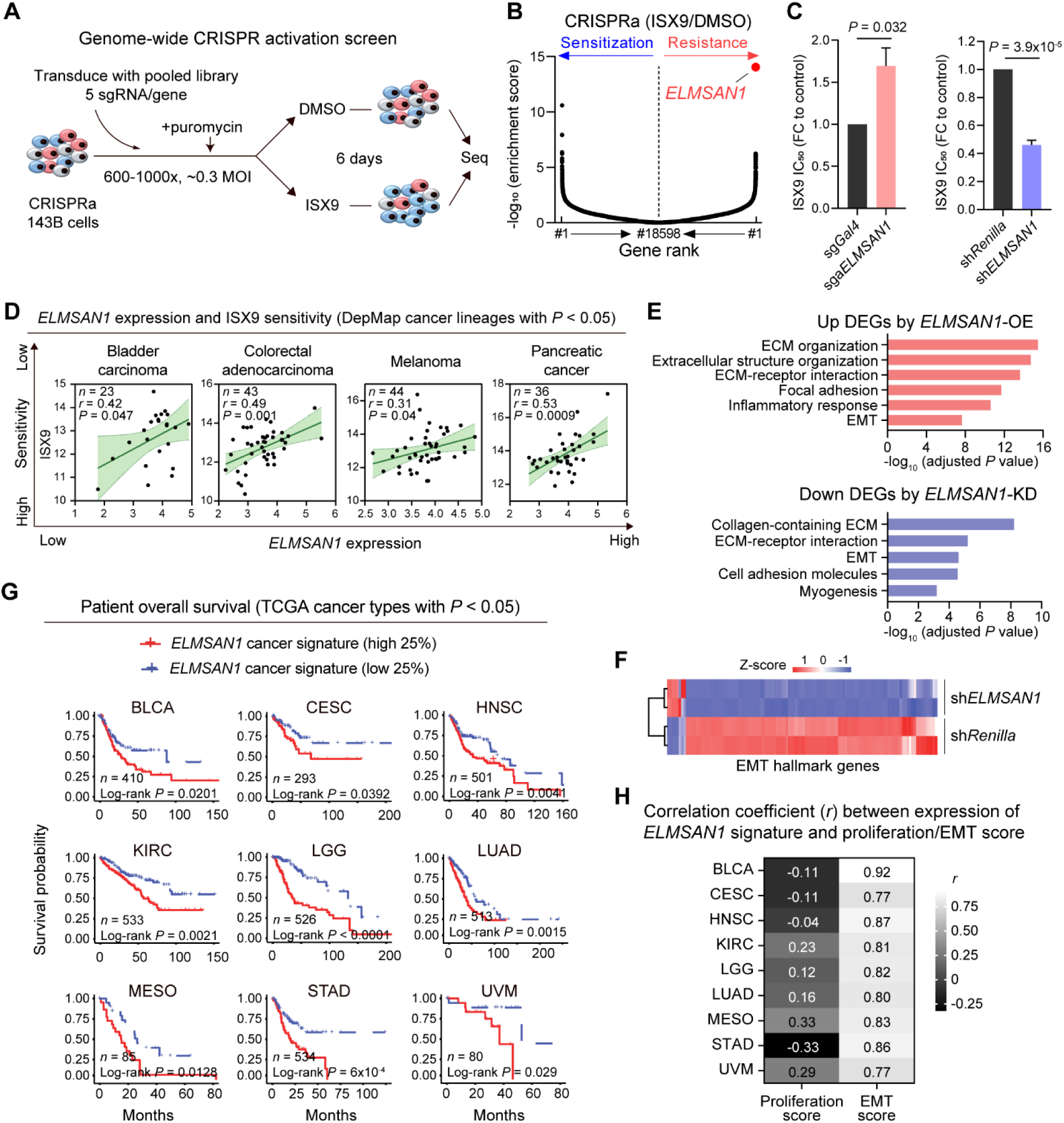
Genome-wide CRISPR activation screen identifies ELMSAN1 as the reprogramming mediator. (A) Schematic of CRISPRa screen. sgRNA: single-guide RNA. MOI: multiplicity of infection. (B) Gene enrichment plot of sensitization and resistance genes ranked by RRA score comparing ISX9 to DMSO in CRISPRa screen. *n* = 2. (C) Relative IC_50_ of ISX9 upon KD and OE of *ELMSAN1*. *n* ≥ 3. (D) Scatterplots depicting the relationship between *ELMSAN1* expression [log_2_ (TPM+1)] (*x*-axis) and sensitivity to ISX9 (area under curve, data from CTD^2^ program) (*y*-axis). All cancer lineages with *P* < 0.05 are shown from DepMap, which are consistently anticorrelated. The shadow area shows 95% confidence intervals. *n* indicates the number of cell lines. (E) Top enriched pathways among upregulated DEGs upon *ELMSAN1*-OE and downregulated DEGs upon *ELMSAN1*-KD. (F) Expression heatmap of EMT hallmark genes among DEGs upon *ELMSAN1*-KD. *n* = 2. (G) Patient overall survival curves stratified by the highest and lowest 25% expression of the *ELMSAN1* cancer signature. All cancer types with *P* < 0.05 are shown from TCGA, which are consistently anticorrelated. *n* indicates patient numbers. (H) Pearson correlation between the expression of *ELMSAN1* cancer signature and proliferation, EMT score (all *P* < 0.0001). Experiments were performed in 143B cells. Data are presented as mean ± SEM. Statistical significance was determined using RRA score (B), two-tailed unpaired t-tests (C), simple linear regression (D), Benjamini-Hochberg method (E), or log-rank (Kaplan-Meier) test (G). See also Figure S2 and Table S3.

ELMSAN1 is a transcriptional corepressor in the mitotic deacetylase complex (MiDAC) comprised of HDAC1/2, ELMSAN1, and DNTTIP1 proteins.^35^ ELMSAN1 binds and activates HDAC1/2,^36^ explaining why OE/KD increased/decreased the IC_50_ of HDACi. *ELMSAN1*-KD downregulated EMT, extracellular matrix, and osteogenic genes (Figures 2E, 2F, and S2I), while *ELMSAN1*-OE upregulated similar genes (Figures 2E and S2J). This is in line with the ISX9 effect to downregulate EMT and cell-of-origin genes (Figure S2K). Thus, ELMSAN1 maintains a mesenchymal state, limiting plasticity and resists the effects of ISX9 and HDACi in osteosarcoma cells. We further explored the clinical importance of *ELMSAN1* in the TCGA cancer database and found a weak correlation between *ELMSAN1* expression and patient survival. We then developed a 14-gene *ELMSAN1* regulatory cancer signature (see STAR Methods). We found that across 32 cancer types in the TCGA database, nine types have a consistently negative relationship (*P* < 0.05) between *ELMSAN1* cancer signature expression and patient overall survival (Figure 2G). Multi-variate analysis was performed to exclude the proliferation index and found strongly positive correlations between the *ELMSAN1* signature and EMT score (*r* > 0.77, all *P* < 0.0001) across all nine cancer types (Figure 2H).

### ELMSAN1 binds to PDC which can be pharmacologically disrupted

ISX9 did not bind to HDAC1/2 proteins (Figure S2B). Both HDAC1 co-IP and ELMSAN1 co-IP assays indicated that ISX9 did not change the ELMSAN1-HDAC1/2 interaction (Figures 3A and S3A). This was further confirmed by unchanged HDAC1/2 enzyme activity in the presence of ISX9 (Figure S3B). We therefore hypothesized that ISX9 might disrupt ELMSAN1 interaction with other proteins. Co-immunoprecipitation (co-IP) of ELMSAN1 followed by mass spectrometry (MS) proteomics (IP-MS) revealed that the binding of DLAT and PDHX, two components in PDC, was lost upon ISX9 treatment (Figure 3A).

**Figure 3.**
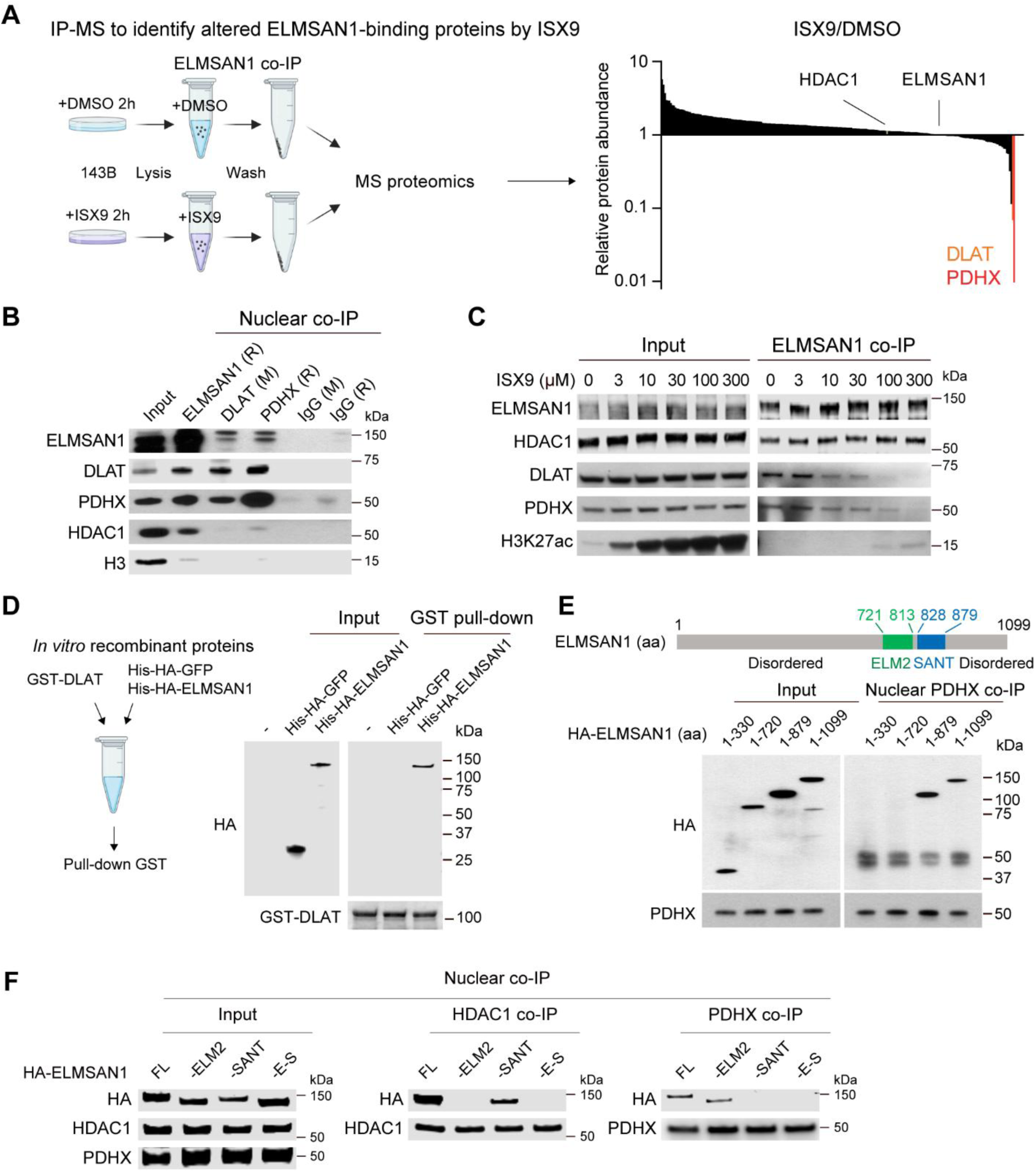
ELMSAN1 binds to PDC which can be pharmacologically disrupted. (A) Schematic of IP-MS experiment and the altered PPI with ELMSAN1 by ISX9 for 2 hours compared to DMSO. ELMSAN1-HDAC1 PPI is not affected by ISX9. (B) Nuclear PPI of ELMSAN1-DLAT/PDHX confirmed by reciprocal co-IP with indicated antibodies in 143B cells. (C) Dose-dependent ELMSAN1-DLAT/PDHX PPI inhibition by ISX9 for 3 hours in 143B cells. (D) Pull-down of recombinant GST-DLAT protein with recombinant His-HA-ELMSAN1 and His-HA-GFP proteins *in vitro*. (E) Nuclear PDHX co-IP with HA-ELMSAN1 truncations expressed in 143B cells to identify critical PPI domains. 721-813aa is the ELM2 domain, and 828-879aa is the SANT domain in the ELMSAN1 protein. (F) Nuclear HDAC1 and PDHX co-IP with HA-ELMSAN1 truncations expressed in 143B cells to identify differential domains required for the ELMSAN1-HDAC1 and ELMSAN1-PDHX binding. See also Figure S3.

DLAT is dihydrolipoamide S-acetyltransferase, the PDC E2 subunit, responsible for generating acetyl-CoA, while PDHX anchors the E3 to the E2 subunit. PDC is vital in linking glycolysis to the tricarboxylic acid cycle in the mitochondrial matrix. It is also present in the nucleus providing acetyl-CoA substrate for histone acetylation.^37,38^ ELMSAN1 is a nuclear protein, and we confirmed the reciprocal ELMSAN1-DLAT/PDHX protein-protein interaction (PPI) (Figures 3B and S3C) and PPI inhibition (Figure S3D) in the nuclei by co-IP of nucleus extracted proteins. The PPI inhibition exhibited an ISX9 dose-dependent manner: 10-30 µM partially and ≥ 100 µM completely inhibited PPI (Figures 3C and S3E). PPI inhibition was also observed in lymphoma and lung cancer cells (Figure S3F), indicating a general mechanism. HDAC1 showed minimal or no interactions with DLAT/PDHX compared with ELMSAN1 (Figure 3B).

In the IP-MS experiment, we observed a modest increase in ELMSAN1 PPI with several proteins (< 5-fold) upon ISX9 treatment (Figure 3A), in contrast to the substantial decrease in binding with DLAT (∼20-fold) and PDHX (> 100-fold). Among the proteins with increased binding, two had relatively high abundance: H2AC21 and H2AZ1, belonging to canonical and variant histone H2A, respectively. The increased binding was validated by ELMSAN1 co-IP and western blot (Figure S3E). The modest increase in binding could suggest that the proximity of ELMSAN1 to chromatin is enhanced, potentially facilitating its transcriptional regulation. However, knocking down H2AZ1 and H2AC21 did not alter the ISX9-induced reduction in cell growth (Figure S3G). These results suggest that the increased binding of ELMSAN1 to these proteins does not mediate the ISX9 effect.

To better study the ELMSAN1-PDC PPI, we generated *in vitro* recombinant Glutathione S-transferase (GST)-DLAT, His-ELMSAN1 proteins and *in vitro* translated His-HA-ELMSAN1, His-HA-GFP proteins. Pull-down experiments showed that DLAT directly interacted with ELMSAN1 (Figures 3D and S3H). Similar to co-IP in cell lysates, ISX9 inhibited ELMSAN1-DLAT PPI in the *in vitro* pull-down assay in a dose-dependent manner (Figure S3H). To define critical ELMSAN1 domains mediating the interaction, we expressed HA-tagged ELMSAN1 truncated proteins followed by co-IP of PDHX and DLAT in 143B cell nuclei. The 1-330aa and 1-720aa truncations were not pulled down by nuclear PDHX and DLAT, but the 1-879aa truncation and full-length (1-1099aa) ELMSAN1 were (Figures 3E and S3I-S3K). This demonstrates that the nPDC-ELMSAN1 interaction requires 721-879aa, the ELM2-SANT domain. To better define the interacting domain, we deleted ELM2, SANT, and ELM2-SANT domains from HA-tagged ELMSAN1 protein followed by nuclear co-IP of PDHX and HDAC1. Notably, we found a different usage of domains for interaction: the SANT domain deletion disrupted ELMSAN1-PDHX binding, whereas the ELM2 domain deletion disrupted ELMSAN1-HDAC1 binding (Figures 3F and S3L). The utilization of the ELM2 domain in the ELMSAN1-HDAC1 interaction aligns with its reported role in mediating interactions between other ELM2-SANT domain-containing proteins and HDAC1/2.^39^ We used protein and small molecule docking analyses to confirm that ISX9 was well accommodated into the pocket of the ELMSAN1-DLAT complex through the SANT domain (Figure S3M). Together, these data indicate that the SANT domain (828-879aa) is critical for ELMSAN1-PDC binding. This is in contrast to the ELM2 domain requirement for HDAC1/2 binding, thereby explaining why ISX9 alters PDC activity but not HDAC.

### Inhibiting ELMSAN1-PDC interaction de-represses nucleus-restricted PDC

We next assessed whether nPDC activity was altered upon ELMSAN1-PDC dissociation. We used a PDC enzyme activity assay that immunocaptured native PDC on a dipstick, followed by the visualization of PDC-dependent NADH production. Notably, ISX9 treatment led to higher PDC enzyme activity in the nuclei (Figures 4A, 4B, and S4A-S4D) without changing protein levels of nuclear DLAT, PDHX (Figure S3D) and captured PDC on the dipstick (Figure 4B). PDC activity was not changed in the non-nuclear fraction, including mitochondria and cytosol (Figures 4A, 4B, and S4D), in line with the nuclear localization of the ELMSAN1 protein. Consistently, *ELMSAN1*-KD enhanced nuclear, but not non-nuclear, PDC activity; *ELMSAN1*-OE reduced nuclear, but not non-nuclear, PDC activity (Figures 4A and S4D). nPDC activity was also enhanced by ISX9 in melanoma and breast cancer cells (Figure S4E). As a result, nuclear acetyl-CoA levels were significantly elevated (Figures 4C and S4F) without affecting nuclear CoA levels (Figure S4G). Consequently, the global histone acetylation, including H3K27ac levels (Figure 3C), were significantly elevated. These data indicate that ELMSAN1 acts as a negative regulator of nPDC that can be activated by ISX9 disrupting the inhibitory action of ELMSAN1.

**Figure 4.**
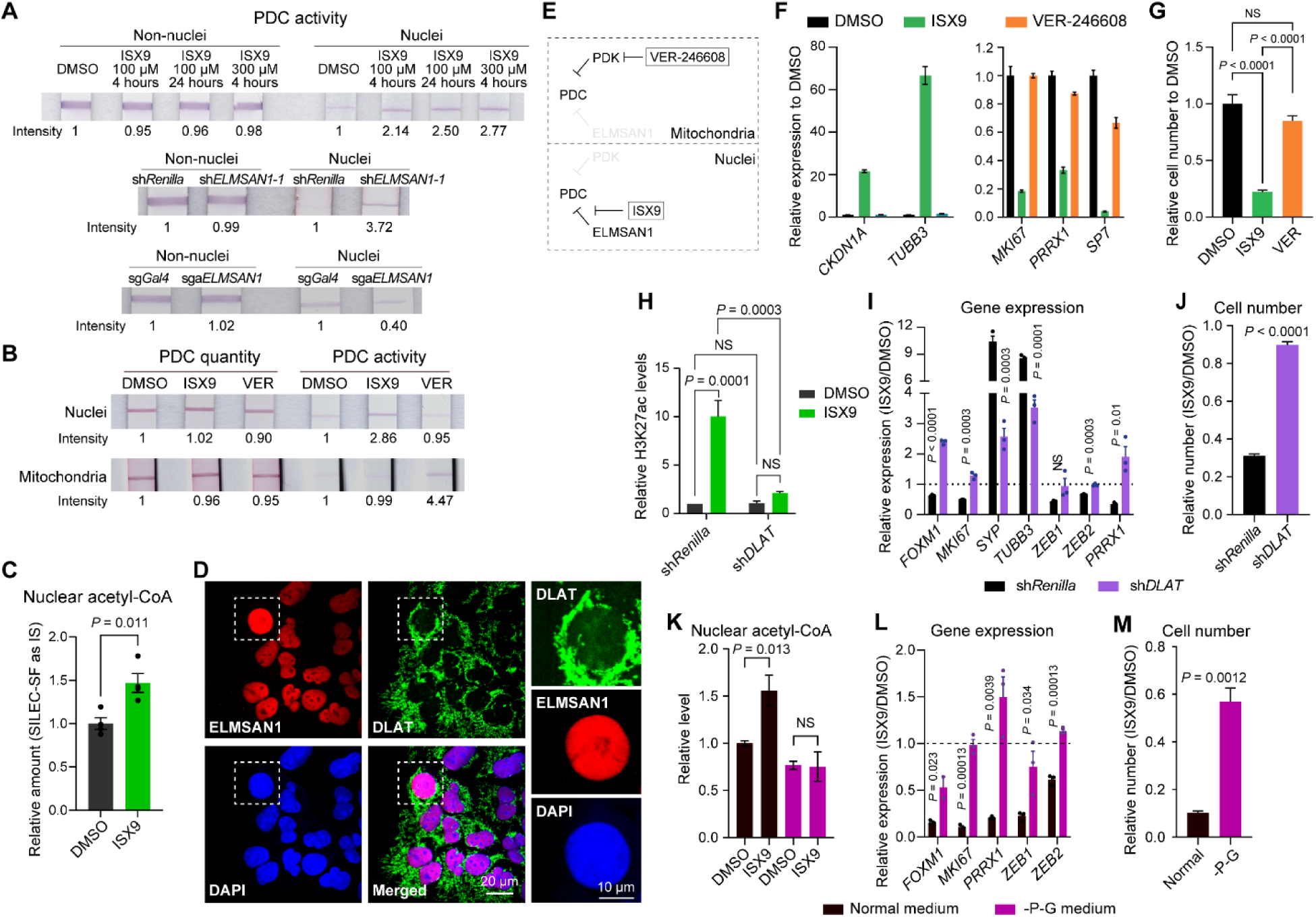
Inhibiting ELMSAN1-PDC interaction de-represses nucleus-restricted PDC. (A) PDC activity assay in indicated conditions with the same cell number. The reaction time between samples in the same image/experiment was the same but was different in different images/experiments. The band intensity was measured. (B) PDC quantity and activity in mitochondria and nuclei under indicated conditions for 3 hours with the same cell number. (C) LC-MS analysis of nuclear acetyl-CoA levels after treated with DMSO and 100 µM ISX9 with the same cell number. Stable isotope labeling of essential nutrients in cell culture-subcellular fractionation (SILEC-SF) was used as an internal standard (IS) control.^50^ *n* = 4. (D) Immunofluorescence images of co-staining ELMSAN1 and DLAT. The higher magnification images of the highlighted field are shown on the right. DLAT is present throughout the nuclei with lesser intensity than the signaling out of the nuclei. Scale bar, 10 µm or 20 µm. (E) Model of organelle-specific regulation of PDC. VER-246608 (20 µM) is a PDK inhibitor affecting mitochondrial PDC. ISX9 (100 µM) is an ELMSAN1-PDC PPI inhibitor affecting nPDC. (F, G) qPCR analysis of MNLT gene expression (F) and relative cell viability (G) under indicated conditions for 3 days. *n* = 3. (H-J) Relative WB signaling of H3K27ac (3 hours, H), qPCR analysis of MNLT gene expression (3 days, I), and cell number (3 days, J) under indicated conditions. *n* = 3. (K-M) LC-MS analysis of nuclear acetyl-CoA levels (3 hours, *n* ≥ 3, K), qPCR analysis of relative gene expression (3 days, *n* = 3, L), and relative cell number (3 days, *n* = 3, M) under DMSO and ISX9 treatment in normal and pyruvate-limiting medium. Data are presented as mean ± SEM. Experiments were performed in 143B cells or PDXC-OS052. Statistical significance was determined using two-tailed unpaired t-tests (C, I, J, L, M) or one-way analysis of variance (ANOVA) with Tukey’s multiple comparison analysis (G, H, K). See also Figure S4.

To better study whether the altered PDC activity is nucleus-restricted, we first performed immunofluorescence staining to show that mitochondria were completely absent and nuclei were intact in the isolated nuclei samples (Figure S4H), confirming a high purity of nuclei for downstream assays. We also co-stained ELMSAN1 and DLAT proteins. As anticipated, DLAT was primarily in mitochondria and present throughout the nucleus with less abundance. ELMSAN1 was confirmed to localize in the nucleus with no overlap with DLAT outside the nucleus (Figure 4D).

Pyruvate dehydrogenase kinase (PDK) is an inhibitory regulator of PDC in mitochondria and it is low/absent in the nucleus (Figure S4I).^37^ Therefore, inhibiting PDK should modulate mitochondrial but not nuclear PDC activity (Figure 4E). Indeed, we observed enhanced PDC activity in mitochondria but no change in the nucleus with the PDK inhibitor VER-246608 (Figure 4B). In contrast, ISX9 increased PDC activity in the nucleus but not in mitochondria (Figure 4B). Both compounds did not change the dipstick-captured PDC quantity (Figure 4B). Correspondingly, VER-246608 had a very limited effect on histone acetylation (Figure S4J), MNLT gene expression (Figure 4F), and cell growth (Figure 4G). Together, multiple lines of evidence demonstrated that the modulation and effect of PDC activity by ISX9 and ELMSAN1 is nucleus-restricted.

Nuclear acetyl-CoA can be generated by different enzymes, including PDC, ACSS2, and ACLY; we asked whether PDC was required in the altered cell state by ISX9. First, we knocked down *DLAT* and found that the ISX9-induced H3K27ac (Figures 4H and S4K), MNLT phenotype (Figure 4I), and cell growth inhibition (Figure 4J) were largely abolished, indicating that ISX9 exerts its function through PDC.

Second, pyruvate is the substrate for PDC to generate acetyl-CoA. We removed pyruvate and its source, glucose, in the medium (-P-G medium) to limit pyruvate for cells (Figure S4L). As expected, the pyruvate-limiting condition reduced nuclear acetyl-CoA levels (Figure 4K). Notably, ISX9 could not increase nuclear acetyl-CoA levels in the pyruvate-limiting condition (Figure 4K). Consequently, the ISX9-induced changes in H3K27ac (Figure S4M), MNLT signature (Figure 4L), and cell growth (Figure 4M) were largely abrogated. The pyruvate-starvation condition growth changes were not attributable to reduced baseline cell growth (Figure S4N). While the H3K27ac activation by ISX9 was pyruvate concentration-dependent (Figure S4O), once a saturating concentration of pyruvate was reached, further increases did not enhance/change the ISX9 effect (Figure S4P). Providing other acetyl-CoA sources such as acetate and acetylcarnitine could not rescue the pyruvate starvation phenotype with ISX9 (Figure S4Q), highlighting a PDC-dependent mechanism. Supplementing lactate in -P-G medium also did not rescue the ISX9 induced phenotype (Figure S4Q), likely due to a primary localization of lactate dehydrogenase in the cytoplasm, again suggesting a nuclear-restricted mechanism. The pyruvate-limiting condition is associated with other processes, such as glycolysis, citrate metabolism, mTOR, and ROS. Nevertheless, the correlation between ablated nuclear acetyl-CoA, H3K27ac, and cell state change indicates that pyruvate plays a vital role in the ISX9 effect. These data demonstrate the essential role of PDC in the ISX9-induced phenotype.

### Derepressing nPDC synergizes with HDAC inhibition to reprogram cancer cells

To profile the altered histone acetylation at the molecular level, we performed CUT&Tag of H3K27ac after 3 hours of ISX9 treatment in PDXCs (Table S4). As expected, we found more increased H3K27ac peaks than decreased (Figures 5A, 5B, and S5A). Similar results were observed in DEG by RNA-seq and differentially accessible regions by ATAC-seq (Figure S5A). We found that most of the increased ATAC peaks had increased H3K27ac levels and most of the increased H3K27ac peaks had an increased chromatin accessibility (Figures 5C, 5D and S5B), consistent with the role of histone acetylation in open chromatin. The H3K27ac pattern change corresponded to gene expression changes: increased H3K27ac peaks were most enriched in genes associated with neuronal and axonal generation, while decreased H3K27ac peaks were most enriched in ribosomal components, Myc targets, E2F targets, and G2-M checkpoint (Figures 5A, 5B and 5D).

**Figure 5.**
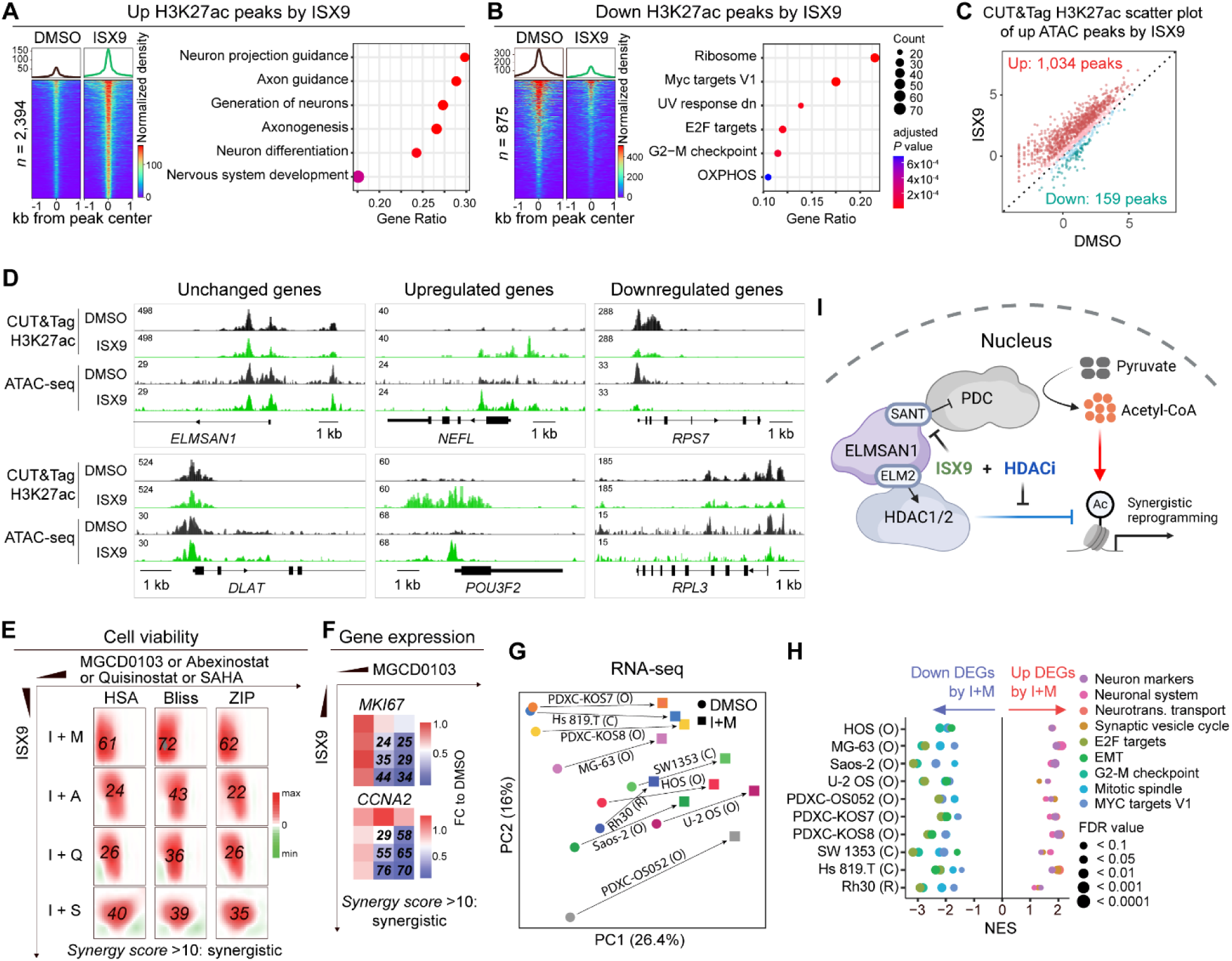
Derepressing nPDC synergizes with HDAC inhibition to reprogram cancer cells. (A, B) Heatmaps and average profiles (left) and enrichment analysis (right) of increased (A) and decreased (B) H3K27ac peaks by 100 µM ISX9 detected by CUT&Tag. *n* = 2. (D) Scatter plot of H3K27ac levels (log_2_RPKM) at the ATAC-seq peaks increased by ISX9. *n* = 2. (E) Genome tracks of representative genes showing H3K27ac and ATAC-seq peaks. *n* = 2 tracks overlaid. (F) Synergy score (italic numbers) of viability under 10×6 dose matrix treatment of ISX9 and four HDACi in PDXC-OS152 calculated by three different models. *n* = 3. (G) qPCR analysis of proliferation gene expression (color) and ZIP synergy score (italic numbers) under 4×3 dose matrix of ISX9 and MGCD0103. *n* = 3. (G, H) Principal component analysis (PCA) (G) and GSEA (H) under I+M and DMSO treatment in 10 sarcoma cell lines including osteosarcoma (HOS, MG-63, Saos-2, U-2OS, PDXC-OS052, PDXC-KOS7, PDXC-KOS8), chondrosarcoma (SW1353, Hs819.T), and rhabdomyosarcoma (Rh30). (I) “Accelerator and brake” model of the mechanism of action of synergistic effect between ISX9 and HDACi in reprogramming cancer cell state. The ELM2 and SANT domains on ELMSAN1 mediate ELMSAN1-HDAC1/2 and ELMSAN1-PDC interaction, respectively. Created by BioRender. Data are presented as mean (F). Statistical significance was determined using the Benjamini-Hochberg method (A, B, H). See also Figure S5.

Because ISX9 increases histone acetylation by providing more acetyl-CoA substrate and HDACi decreased histone acetylation erasure, we wondered if ISX9 and HDACi could lead to a similar phenotype. We used MGCD0103 at 1 µM to inhibit HDAC1/2 activity by 85% (Figure S3B). We found a more rapid activation of H3K27ac by ISX9 (Figures S5C and S5D). PCA based on H3K27ac CUT&Tag levels suggested different changes induced by ISX9 and MGCD0103 (Figure S5E). Analysis of H3K27ac marks and expression of key MNLT genes suggested that ISX9 exerted a distinct or stronger effect than MGCD0103 (Figures S5F-S5I). When treating and withdrawing the compounds on day 6, ISX9-treated PDXCs had persistently suppressed growth for 12 days, in contrast to the fully restored growth of cells receiving MGCD0103 (Figure S5J). In sum, although both HDACi and ISX9 enhance histone acetylation, they exert their effect by different mechanisms. Nuclear acetyl-CoA generation through ISX9 derepressing nPDC provides a local, rapid and genomic loci-specific supply for histone acetylation, thereby reprogramming cell state differently than HDACi treatment.

Next, we assessed whether ISX9 and HDACi have a cooperative effect. Three different models (Bliss, HSA, and ZIP)^41^ consistently indicated synergistic effects between ISX9 and MGCD0103 (I+M) in inhibiting PDXC viability (Figure 5E). The ZIP model demonstrated a synergistic effect between I+M in changing the expression of proliferation genes (Figure 5F). Moreover, ISX9 also synergized with FDA-approved HDACi SAHA (S) and second-generation HDACi Abexinostat (A) and Quisinostat (Q) (Figure 5E). Mechanistically, we treated PDXCs with ISX9, MGCD0103, and I+M at the concentrations that exhibited high synergy scores and performed H3K27ac CUT&Tag (Table S4). In line with synergy, 70-90% of the changed peaks by I+M were exclusive to the combination but not by ISX9 or MGCD0103 alone (Figure S5K). The exclusively downregulated peaks were highly enriched in proliferation genes and Myc targets (Figure S5K), explaining the synergistic effect of I+M in inhibiting cell growth. Interestingly, at the concentration of ISX9 exhibiting high synergy (10-20 µM), ISX9 partially inhibited ELMSAN1-PDC interaction.

The strong synergistic effect of I+M prompted us to examine if it could be applied beyond osteosarcoma. I+M induced a similar transcriptional change in 10 sarcoma cell lines (Figure 5G) with the downregulation of proliferation and mesenchymal genes and the upregulation of neuronal genes and *P21* (Figures 5H and S5L). In rhabdomyosarcoma cells, I+M downregulated the myogenic program transcriptionally and epigenetically and increased accessible regions associated with neuronal genes (Figures S5M-S5P). In other cancer types, I+M again markedly downregulated proliferation genes and the cell-of-origin TF of the corresponding lineages: *SOX10* in melanoma and *FOXA1* in lung, prostate, and breast cancers and mildly activated neuronal markers (Figure S5Q). Analysis of 1,096 cancer cell lines in the pan-cancer DepMap database^42^ suggested that cell-of-origin/lineage-specific TFs are essential genes for corresponding cancer cell survival (Figure S5R), supporting the general benefit of turning off the cell-of-origin signature. Thus, derepressing nPDC synergizes with HDACi to reprogram multiple cancer types. We propose an “accelerator and brake” model. Acetyl-CoA serves as an accelerator; it is induced by ISX9 derepression of nPDC. Inhibition of deacetylation by HDACi releases the countervailing brake. ISX9 and HDACi synergistically alter the H3K27ac landscape and reprogram cell state (Figure 5I).

### Therapeutic cancer cell reprogramming *in vivo*

To test the pharmacological efficacy of small molecules *in vivo*, we employed four PDX models of osteosarcoma differing in mutations (including *TP53*, *PTEN*, *MYC*, *AKT1*, *CCNE1*), disease stage at sample acquisition (diagnostic or metastatic), and treatment status (pre- or post-treatment), two PDX models of Ewing sarcoma (EWS) (Figure S6A),^43^ and one orthotopic xenograft model of rhabdomyosarcoma (RMS) by administrating the combination of ISX9-LD (25-40 mg/kg, once daily) and MGCD0103. An evident and consistent inhibition of tumor growth by I+M was observed in all models (Figures 6A and S6B-S6H). ISX9-LD and MGCD0103 exhibited a cooperative effect in reducing tumor growth *in vivo* (Figure S6E). Notably, all three post-treatment tumor-derived PDX models (PDX-OS052, OS152, and OS525) responded to I+M treatment (all *P* < 0.001). Given that these models represent therapy-resistant tumors, *in vivo* epigenetic reprogramming to alter clinically relevant outcomes is of considerable interest.

**Figure 6.**
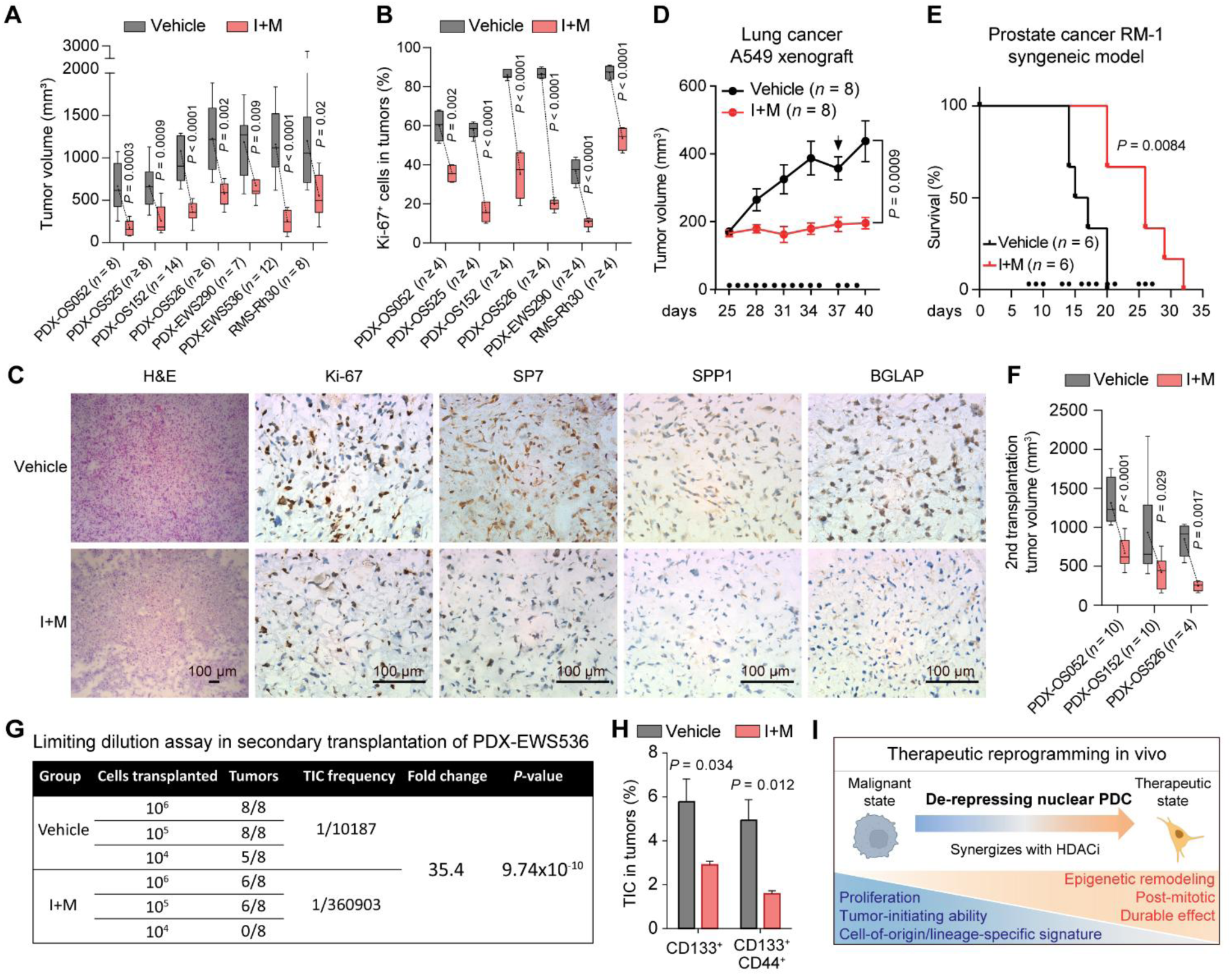
Therapeutic cancer cell reprogramming *in vivo*. (A) Tumor volumes under vehicle and I+M treatment in four PDX models of osteosarcoma, two PDX models of Ewing sarcoma and one xenograft model of rhabdomyosarcoma. (B) Quantification of Ki-67 positive cells in tumors under vehicle and I+M treatment in different models. (C) Representative immunohistochemistry images in PDX-OS052 tumors under vehicle and I+M treatment. Scale bar, 100 µm. (D) Tumor growth curve of A549 xenograft under vehicle and I+M treatment. Arrow indicates the euthanasia of a vehicle-treated mouse because of tumor ulceration. (E) Survival curve of RM-1 syngeneic mice under vehicle and I+M treatment determined by a total tumor volume of 800 mm^3^. (F) Tumor volumes in secondary transplantation with vehicle and I+M treated tumors in three PDX models. No treatment in the secondary transplanted mice. (G) Limiting dilution analysis of TIC frequency by vehicle and I+M treatment in PDX-EWS536. Primary transplanted mice were treated with vehicle or I+M for 8 days and then tumors were harvested/transplanted into secondary recipients with indicated cell number without treatment. Tumor frequency was determined at 6 weeks in the secondary transplantation. (H) Flow analysis of TIC (defined by CD133^+^ or CD133^+^CD44^+^) percentage in PDX-OS052 tumors under vehicle and I+M treatment. *n* = 4 tumors. (I) Key features of therapeutic cancer reprogramming induced by derepressing nuclear PDC. Data are presented as box plots divided by median and Tukey-style whiskers (A, B, F) or mean ± SEM (D, H). Sample numbers (*n*) are indicated in each panel. Dots on *x*-axis indicate drug administration timepoints. Statistical significance was determined using two-tailed unpaired t-tests (A, B, D, F, H), or log-rank (Mantel-Cox) test (E). See also Figures S6 and S7.

Immunohistochemistry showed a similar phenotypic change *in vivo*. A significant reduction of Ki-67 and an increase in P21 were observed in tumors in I+M treated mice (Figures 6B, 6C, and S6C). Additionally, immunohistochemistry showed that I+M diminished osteogenic protein expression, including SP7, BGLAP, and SPP1 in PDX tumors (Figures 6C and S6C). Neuronal marker TUBB3 was detected in EWS PDX tumors by I+M (Figure S6G). MYOD1 protein in rhabdomyosarcoma xenograft tumors was downregulated by I+M (Figure S6H). We further tested lung cancer A549 xenograft and prostate cancer RM-1 syngeneic models; I+M again delayed tumor growth and prolonged mouse survival (Figures 6D, 6E, and S6I).

Secondary transplantation studies were conducted to assess the impact of I+M treatment on tumor stem-cell function or tumor-initiating cell (TIC) ability. Secondary tumors from the I+M group grew considerably slower than those from the vehicle group in all tested PDX models despite the absence of further I+M treatment (Figures 6F and S6J). Limiting dilution analysis demonstrated a marked decrease in TIC frequency following I+M treatment in both EWS PDX model (35.4-fold, *P* = 9.74×10^−10^) (Figure 6G) and osteosarcoma PDX model (5.6-fold, *P* = 0.0036) (Figure S6K). Flow cytometry further confirmed the reduction of the TIC population (Figures 6H and S6L).

The capacity of I+M to modulate the cancer cell state raised concern about its effects on normal cells. We grossly evaluated the *in vivo* toxicity by administrating I+M for three weeks in wild-type mice (Figure S7A). Body and vital organ weights (Figures S7B and S7C), complete blood counts (Figure S7D), and mesenchymal stromal cell (MSC) percentage and function were not significantly affected (Figures S7E-S7G). These data suggest limited drug toxicity and a potential therapeutic window.

## Discussion

Nuclear metabolism is increasingly appreciated to be important for epigenetic regulation and cell state control.^44–49^ Although nuclear acetyl-CoA only constitutes a small fraction of the acetyl-CoA pool in a whole cell,^50^ it provides a local and possibly more timely supply of substrate for histone acetylation. Consistent with this notion, nPDC has been reported to regulate important biological processes, including cell cycle, early embryogenesis, tumorigenesis, and transcription.^37,51–53^ However, processes regulating nPDC require further definition as constitutive activation of nPDC has been proposed due to the absence of PDK in the nucleus.^37,47^ Our findings indicate the inhibition of nPDC by ELMSAN1 through direct binding. Importantly, the regulation of PDC by ELMSAN1 is restricted to the nucleus. The marked cell state effects on malignant cells of nPDC regulation highlight the importance of studying metabolism in a subcellular compartmentalized context.

Epigenetic plasticity in cancer is associated with distinct phenotypes and clinical outcomes. Non-genetic adaptive mechanisms following drug therapy result in the generation of drug-tolerant cells, which are quiescent/slowly dividing persister cells.^30,54,55^ These can be the consequence of “transdifferentiation,” acquiring neuroendocrine traits or alternative stem-cell programs in prostate, pancreatic, basal cell carcinomas or melanoma.^11,56–62^ The drug-resistant state is often accompanied by a high level of malignant stemness and eventually leads to treatment failure.^13,54,56,57,60,61^ In contrast, in this study, we exploit epigenetic plasticity to show that a therapeutic cell state transition can be pharmacologically induced in sarcomas to achieve a significant decrease in stemness, cell-of-origin gene expression *in vitro* and durable loss of tumor-initiation ability *in vivo*, even in the context of therapy-resistant disease (Figure 6I). This is the consequence of active epigenetic reprogramming rather than adaptation to survive following therapies. Importantly, similar therapeutic effects were observed in lung and prostate cancers.

Pharmacologic disruption of ELMSAN1-nPDC interaction derepresses nPDC leading to epigenetic remodeling of the histone acetylation landscape. ELMSAN1 is highly conserved in evolution and known to participate in the HDAC complex MiDAC.^35,36,40,63^ Our study reveals the critical role of ELMSAN1 in nuclear pyruvate metabolism that then provides ELMSAN1 with two complementing roles in regulating histone acetylation. Modulating these two functions of ELMSAN1 leads to the synergistic effects of derepressing nPDC and inhibiting HDAC on tumor growth, animal survival, and a durable loss of tumor-initiation ability even after drug removal and therapy-resistant disease. Interestingly, the two roles of ELMSAN1 rely on different domains to exert function: the ELM2 domain mediates ELMSAN1-HDAC interaction and the SANT domain mediates the ELMSAN1-nPDC interaction. In sum, derepressing nPDC rewires the histone acetylation landscape sufficiently and durably to alter the cancer phenotype. The result is an anti-tumor effect across multiple human cancer xenograft models and a depletion of cancer stem cells.

### Limitations of the study

Although ISX9 showed *in vivo* efficacy, we recognize that further optimization of ISX9 or analogs will be needed for clinical translation. Although we defined the critical PPI domain in ELMSAN1, future work to resolve the structural basis for the inhibitory function of ELMSAN1 on PDC will be important. Understanding the spatial localization and generation of acetyl-CoA mediated by ELMSAN1 and the specificity of acetyl-CoA regulation within the nucleus will be challenging and important.

## RESOURCE AVAILABILITY

### Lead contact

Further information and requests for resources and reagents should be directed to and will be fulfilled by the lead contact, David T. Scadden.

### Materials availability

All unique/stable reagents generated in this study are available from the lead contact by request.

### Data and code availability

- Raw and processed RNA-seq, ATAC-seq, CUT&Tag, and CRISPR data from this study have been deposited in the Gene Expression Omnibus and are publicly available as of the date of publication. Accession numbers are listed in the key resources table.
- Raw data of graphs and original western blot images are in Supplemental file Data S1.
- Any additional information required to reanalyze the data reported in this paper is available from the lead contact upon request.

## ACKNOWLEDGEMENTS

We would like to thank all Scadden lab members, Raul Mostoslavsky, Timothy Graubert, Peter Miller, Hongkui Deng for thoughtful discussion and Julia Joung, Wentao Yu, Celia Torres, Wenxiang Zhang, Yu Wang, Xubo Niu, Weikun Xia, Min Yang for technical support; Jacqueline Lees (MIT) for providing OS2674 cells; Bertha Brodin (Karolinska Institute) for providing PDXC-KOS7 and PDXC-KOS8; Ulandt Kim and Zhenya Li at the MGH Next Generation Sequencing core for assistance with the RNA-seq, ATAC-seq, CUT&Tag and gRNA sequencing; Charles Vidoudez at the Harvard Center for Mass Spectrometry for assistance with the metabolite studies, Mei Chen and Steven Kolakowski at the Harvard Center for Mass Spectrometry for assistance with the proteomics studies, MGH Histopathology Research Core, HSCI CRM Flow Cytometry Core, and Animal facility at Center for Comparative Medicine at MGH. This research was supported by two grants from the Sarcoma Foundation of America to D.T.S and T.Z., and the Gerald and Darlene Jordan Professorship to D.T.S.

## AUTHOR CONTRIBUTIONS

Conceptualization, T.Z. and D.T.S.; Methodology, T.Z., L.H., R.I.S., E.A.S.C., and D.T.S.; Investigation, T.Z., L.H., J.X., Y.X., J.G.V.V., M.M., L.C., C.R., T.Fa., T.Fu., M.D., J.A.B.G.; Formal analysis: T.Z., L.P.W., S.M.; Writing – Original Draft, T.Z., D.T.S.; Writing – Review & Editing: all authors; Visualization: T.Z.; Project Administration, S.P.G., Z.D., D.B.S., R.I.S., E.A.S.C.; Funding Acquisition, T.Z. and D.T.S. Resources, L.C.S., E.A.S.C.; Supervision, D.T.S.

