## Supplemental materials and figures for "Interrupting ELMSAN1 repression of nuclear acetyl-CoA production therapeutically reprograms cancer cells"


##### EXPERIMENTAL MODEL AND STUDY PARTICIPANT DETAILS

##### Mice

All animal experiments were performed in accordance with national and institutional guidelines. Mice were housed in the Massachusetts General Hospital (MGH) Animal Research Facility on a 12-hour light/dark cycle with stable temperature (22°C) and humidity (60%). All procedures were approved by MGH Internal Animal Care and Use Committee. Animals were randomly assigned to experimental and control groups with a similar number of mice. When possible, the investigators were blinded during experiments and data analysis.

##### Patient-derived xenograft models

Primary patient-derived xenografts (osteosarcoma: PDX-OS052, PDX-OS152, PDX-OS525, PDX-OS526; Ewing sarcoma: PDX-EWS290, PDX-EWS536) were kindly provided by Dr. Alejandro Sweet-Cordero's group at the University of California, San Francisco. Cells were first transplanted for expansion by subcutaneously injecting into NOD-scid gamma (NSG) (NOD.Cg-Prkdc$^{scid}$ Il2rg$^{tm1Wjl}$, The Jackson Laboratory, 005557) mice with Matrigel (Corning, 354230) 1:1 diluted by PBS. When reaching the maximal size, tumors were resected and cut into small fragments, then digested in 0.25% collagenase type I (Stem cell technologies, 07902) with 2 mg/ml dispase (Thermo Fisher Scientific, 17105041) and 12 U/ml DNase I (Thermo Fisher Scientific, 90083) for 10-20 min at 37°C with agitation (120 rpm). One more digestion of remaining tumor tissues may be applied. After digestion, samples were filtered through a 70 µm filter, and erythrocytes were lysed in ACK lysis buffer (Quality Biological, 118-156-721) for 5 min on ice. Cell number was counted with trypan blue (Corning, 25-900-CI) staining to identify live cells. An equal number of live cells (PDX-OS052: $4x10^6$, PDX-OS152: $2x10^6$, PDX-OS525: $3x10^6$, PDX-OS526: $2x10^6$, PDX-EWS290: $3x10^6$, PDX-EWS536: $8x10^6$) were subcutaneously transplanted into flank of NSG mice. When the tumor reached a mean volume of approximately 100-200 $mm^3$, mice were randomly distributed into different groups. For ISX9 single-agent treatment (ISX9-HD), it was intraperitoneally injected at 50 mg/kg, twice daily. For I+M combination treatment, ISX9 was intraperitoneally injected at 25-40 mg/kg, once daily (ISX9-LD); MGCD0103 was intraperitoneally injected at 15-25 mg/kg once daily. ISX9 (Selleckchem, S7914) was dissolved at a concentration of 2 mg/ml in 20% (2-hydroxypropyl)-β-cyclodextrin (HBC, MilliporeSigma, 332607). MGCD0103 (MedChemExpress, HY-12323) was dissolved in 0.1 N HCl (Fisher Scientific, SA49) in PBS. The same volume of vehicle was used as control. Body weights were monitored every day or every other day. We observed mild weight loss in I+M treated NSG mice. This is likely due to the different dose tolerance in NSG mice because such weight loss did not happen in wild-type mice. If the body weight loss was >10%, drug administration was suspended until weight became stable. Mice were generally kept in healthy condition. Tumor volumes were measured twice a week using SPI electronic calipers (MSC, 03883980) and calculated using the formula $v = l \times w^2/2$, where l means length and w means width. Mice with a tumor burden of 1,500 $mm^3$, tumor ulceration/bleeding, sign of infections, respiratory distress, impaired mobility, disability to walk and access food, leg paralysis, >20% reduced in body weight or general cachexia were humanely euthanized. Tumors were harvested at the endpoint for histological analysis, cell sorting, and secondary transplantation.

**Xenograft models**

Rhabdomyosarcoma Rh30 cells ($2x10^6$) were intramuscularly transplanted into NSG mice with Matrigel 1:1 diluted by PBS. Drug treatment and tumor monitoring were described above.

**Syngeneic model**

Mouse prostate cancer cell lines RM-1 ($5x10^6$) were subcutaneously transplanted into C57BL/6J mice with Matrigel 1:1 diluted by PBS. Drug treatment and tumor monitoring were described above.

##### Cell culture

143B, HOS, MG-63, Saos-2, U-2 OS, OS2674, PDXC-KOS7, PDXC-KOS8, PDXC-OS052, PDXC-OS152, PDXC-OS526, SW1353, Hs 819.T were cultured in DMEM (Thermo Fisher Scientific, 11995-065) with 10% fetal bovine serum (FBS, Thermo Fisher Scientific, A3160501), 1% Penicillin/Streptomycin (PS, Thermo Fisher Scientific, 15140122) and 1% Glutamine (Corning, 25-005-CI). Rh30 were cultured in IMDM (Thermo Fisher Scientific, 12440053) with 20% FBS, 1% PS, 1% Glutamine, and 1% Insulin-Transferrin-Selenium (Thermo Fisher Scientific, 41400045). A549, MDA-MB-231, and MCF7, BT549, SK-MEL-28, A-375, COLO 679, PC-3 were cultured in RPMI (Thermo Fisher Scientific, 11875119) with 10% FBS, 1% PS, and 1% Glutamine. For pyruvate-limiting culture, cells were cultured in no glucose, no pyruvate medium (Thermo Fisher Scientific, 11966025) supplemented with 10% FBS, 1% PS, and 1% Glutamine. ISX9, MGCD0103, VER-246608, and other chemicals were used as indicated; their detailed information can be found in the Key Resource Table. Cells were split every 3-6 days, depending on the confluency. All cells were maintained at 37°C with 5% $CO_2$. PDXC-OS052, PDXC-OS152, PDXC-OS526, and PDXC-OS384 were kindly provided by Dr. Alejandro Sweet-Cordero's lab at the University of California, San Francisco. PDXC-KOS7 and PDXC-KOS8 were kindly provided by Dr. Bertha Brodin's lab at Karolinska Institute. OS2674 was kindly provided by Dr. Jacqueline Lees's lab at Massachusetts Institute of Technology. Rh30 was obtained from the Childhood Cancer Repository. 143B, HOS, MG-63, Saos-2, U-2 OS, SW1353, Hs 819.T, A549, MDA-MB-231, and MCF7, BT549, SK-MEL-28, A-375, and PC-3 were from the American Type Culture Collection.

##### Screening platform for Stemness and Viability Inhibitors

We used OS2674 cells for the screen, which was derived from osteosarcoma GEMM. OS2674 cells were seeded into the black wall, clear-bottom 96-well plate (Corning, 3904) at a density of 10,000 cells per well in culture medium. Edge wells were excluded. The next day, the medium was changed to MEMα with 15% FBS, 1% PS and 1% Glutamine, and 10 mM β-glycerophosphate (MilliporeSigma, G9422). We found that cells exhibited better mineralization in MEMa than DMEM medium. We screened the bioactive compound library (3,046 chemical compounds including FDA-approved and clinical trial drugs) from Selleckchem (L1700, 2018 version). They were diluted serially in MEMα and added into the wells individually at the final concentration of 0.5 µM and 5 µM. DMSO was used as a negative control. > 10 wells of DMSO were used, whose average of PB and ARS signaling was used for calculation. Four days after induction, cell viability and calcium deposits were measured sequentially. First, cells were stained by directly adding 10x PrestoBlue viability reagent (Thermo Fisher Scientific, P50200) into the medium. After 1 hour of incubation, fluorescence was measured at 560/590 nm by a SpectraMax i3x Multi-Mode Microplate Reader. Then, cells were washed with PBS and stained with Alizarin Red S (ARS) stain solution (MilliporeSigma, TMS-008-C) 40 µl/well at room temperature for 15 min. After $H_2O$ wash twice with gentle rocking, 100 µl/well 10% acetic acid was added to extract the calcified mineral for 5 min and then transferred to new plates for a colorimetric quantification at 405 nm by plate reader. Medium without cells was used as a background control for PrestoBlue and ARS staining. We calculated the results by using raw data as below:

$$\text{Fold change of ARS} = \frac{\text{screening well} - \text{background well}}{\text{DMSO well} - \text{background well}}$$

$$\text{Percentage of viability} = \frac{\text{screening well} - \text{background well}}{\text{DMSO well} - \text{background well}} \text{ x } 100\%$$

**Cell viability, $IC_{50}$ and synergy calculation**

Cells were seeded into 384-well or 96-well plates. The next day, small molecules were serially diluted in medium and added to cells either alone or in a dose matrix. Prestoblue staining or CellTiter-Glo® Luminescent Cell Viability Assay (Promega, G7571) was performed according to the manufacturer's protocol. $IC_{50}$ was calculated by GraphPad Prism. SynergyFinder[41] was used to calculate the score of synergy by using highest single agent (HSA), Bliss, and Zero interaction potency (ZIP) models.

**Senescence, HDAC activity, and pyruvate-Glo assay**β-galactosidase staining was performed using the senescence β-galactosidase staining kit (CST, 9860S). Fibroblasts were used as a positive control, which was prepared by treating with 10 µg/ml mitomycin C (MilliporeSigma, M4287) for 3 hours and further culturing for several days. HDAC activity assay was performed using recombinant HDAC1 (BPS Bioscience, 50051) and recombinant HDAC2 (BPS Bioscience,

50052) with HDAC activity assay kit (Abcam, ab1432). 0.2 μg HDAC1 and 1 μg HDAC2 protein was used per reaction (well). Intracellular pyruvate was measured by pyruvate-Glo assay kit (Promega, J4051) after culturing cells in -P-G medium for indicated time. The same number of cells were used for each timepoint.

**Reverse transcription-quantitative PCR**

Total RNA was extracted using Direct-zol RNA MiniPrep Kit (Zymo Research, R2052). cDNA was synthesized from total RNA using iScript cDNA Synthesis Kit (Bio-Rad, 1708891). qPCR was performed using iTaq Universal SYBR Green Supermix (Bio-Rad, 1725121) on a CFX384 Real-Time System (Bio-Rad). The data were analyzed using the $2^{-\Delta\Delta Ct}$ method. ACTB was used as a housekeeping gene. The following primers were used for qPCR analysis:

| Gene | Forward (5' -> 3') | Reverse (5' -> 3') |
|---|---|---|
| *ACTB* | AGAGCTACGAGCTGCCTGAC | AGCACTGTGTTGGCGTACAG |
| *MKI67* | GAAAGAGTGGCAACCTGCCTTC | GCACCAAGTTTTACTACATCTGCC |
| *CCNA2* | CTCTACACAGTCACGGGACAAAG | CTGTGGTGCTTTGAGGTAGGTC |
| *FOXM1* | TCTGCCAATGGCAAGGTCTCCT | CTGGATTCGGTCGTTTCTGCTG |
| *P21* | AGGTGGACCTGGAGACTCTCAG | TCCTCTTGGAGAAGATCAGCCG |
| *RUNX2* | TGGTTACTGTCATGGCGGGTA | TCTCAGATCGTTGAACCTTGCTA |
| *SP7* | CCTCTGCGGGACTCAACAAC | AGCCCATTAGTGCTTGTAAAGG |
| *SPP1* | GAAGTTTCGCAGACCTGACAT | GTATGCACCATTCAACTCCTCG |
| *IBSP* | TTCATTGAATGGTTTGAGGTTG | AGTGTTGCATAGGTAGTGCGATT |
| *COL1A1* | GAGGGCCAAGACGAAGACATC | CAGATCACGTCATCGCACAAC |
| *ALPL* | CCTGCCTTACTAACTCCTTAGTGC | CGTTGGTGTTGAGCTTCTGA |
| *NEUROD1* | GGTGCCTTGCTATTCTAAGACGC | GCAAAGCGTCTGAACGAAGGAG |
| *TUBB3* | TCAGCGTCTACTACAACGAGGC | GCCTGAAGAGATGTCCAAAGGC |
| *SYP* | TCGGCTTTGTGAAGGTGCTGCA | TCACTCTCGGTCTTGTTGGCAC |
| *SYN1* | CGATGCCAAATATGACGTGCGTG | AGCATCGCAGAGCCAGTATTGG |

| | | |
|---|---|---|
| *MAP2* | AGGCTGTAGCAGTCCTGAAAGG | CTTCCTCCACTGTGACAGTCTG |
| *PRRX1* | TGCAGGCTTTGGAGCGTGTCTT | CTCATTCCTGCGGAACTTGGCT |
| *SNAI2* | ATCTGCGGCAAGGCGTTTTCCA | GAGCCCTCAGATTTGACCTGTC |
| *ZEB1* | GGCATACACCTACTCAACTACGG | TGGGCGGTGTAGAATCAGAGTC |
| *ZEB2* | AATGCACAGAGTGTGGCAAGGC | CTGCTGATGTGCGAACTGTAGG |
| *SOX10* | ATGAACGCCTTCATGGTGTGGG | CGCTTGTCACTTTCGTTCAGCAG |
| *FOXA1* | GCAATACTCGCCTTACGGCTCT | GGGTCTGGAATACACACCTTGG |
| *ELMSAN1* | GCACAAGCCATCAGTCATCGTC | CTGCTTTGGTTTCCGCACGGAA |

**Lentivirus for knockdown and overexpression**

Lentiviral constructs were transfected with VSVG and Δ8.9 to HEK293T cells using FuGENE6 (Promega, E2692). Viral supernatant was centrifuged and concentrated using Lenti-X Concentrator (TaKaRa, 631232). The concentrated virus was transduced into recipient cells supplemented with 8 μg/ml polybrene. The infected cells were then selected with puromycin 2 days after transduction.

Knockdown experiments employed Tet-pLKO-puro vector (Addgene, #21915) or pLV-puro-U6 vector. 1 μg/ml doxycycline (DOX) was used to induce knockdown. The following shRNAs were used:

sh*ELMSAN1*-1: CTACTTCAATGCCATCATATC.

sh*ELMSAN1*-2: CTTTCCACGCAGCCAAGAAAC.

sh*DLAT*: CAACCGAAGTAACAGATTTAA.

sh*H2AC21*: GAGTACCTGACCGCGGAAATT.

sh*H2AZ1*: ATTCGAGTCTTAACCATATTT.

*ELMSAN1* overexpression was performed by using CRISPRa with sgRNA protospacer sequence (GGCGACCGACTCGTGAGTGG) or using pLV-Puro-CMV>hMIDEAS [NM_001367710.1].

**Immunohistochemistry**

Tumors were dissected, cut, and embedded in optimal cutting temperature (OCT) compound (Tissue-Tek, 4583) and sectioned at 5 μm, followed by staining with hematoxylin and eosin (H&E), anti-SP7 (Novus Biologicals, NBP3-10930, 1:200), anti-SPP1 (OPN) (Abcam, ab8448, 1:200), anti-BGLAP (OCN) (Abcam, ab93876, 1:500), anti-TUBB3 (TUJ1) (BioLegend, 801201, 1:200), anti-NEFL (CST, 2837T, 1:100), anti-MYOD1 (Abcam, ab133627, 1:500), anti-P21 (CST, 2947S, 1:50), and anti-Ki67 (Thermo Fisher Scientific, 14-5698-82, 1:100). For quantification of Ki-67, P21 and MYOD1 positivity, > 4 random fields from multiple sections were analyzed.

**Immunofluorescence**

Cells were seeded and treated on round glass cover slides coated with 0.1% gelatin (MilliporeSigma, G9136). After treatment, cells were first fixed in 4% paraformaldehyde at room temperature for 20 min and washed by PBS three times. Then, cells were permeabilized with 0.1% Triton-X100 (MilliporeSigma, T8787) and blocked with blocking buffer (PBS containing 0.1% Tween 20 and 2.5% donkey serum) at 37°C for 1 hour. Primary antibodies incubation was performed at 4°C overnight in blocking buffer with antibodies: anti-TUBB3 (CST, 5568, 1:500), anti-Ki67 (Thermo Fisher Scientific, 14-5698-82, 1:100), anti-SP7 (Novus Biologicals, MAB7547, 1:200), anti-ELMSAN1 (Thermo Fisher Scientific, PA5-51825, 1:100), anti-DLAT (CST, 12362, 1:100), anti-PDK1 (Proteintech, 18262-1-AP, 1:200). The next day, cells were washed with PBS three times and probed with secondary antibodies (Jackson ImmunoResearch) at 37°C for 1 hour in blocking buffer. Cells were then washed with PBS three times and counterstained with 1 μg/ml DAPI (MilliporeSigma, 10236276001). For mitochondrial and nuclear staining during nuclei isolation, the cells or nuclei from each step were cytospined to the slides and stained with 50 nM MitoTracker Green (MedChemExpress, HY-135056) and 5 μM DRAQ5 (Thermo Fisher Scientific, 62251) in DMEM for 30 min at room temperature. Finally, slides were mounted by ProLong Diamond Antifade Mountant (Thermo Fisher Scientific, P36965) on microscope slides and cells were imaged by Leica SP8 confocal microscope. Quantification of the percentage of positive cells was done under identical microscopy settings between samples. 3-6 randomly selected fields were analyzed.

**Co-immunoprecipitation**

For whole cell lysate co-IP, cells were trypsinized and lysed in buffer containing 20 mM Tris, pH 7.5, 137 mM NaCl, 1 mM $MgCl_2$, 1 mM $CaCl_2$, and 1% NP-40, supplemented with 2 units/mL benzonase (MilliporeSigma, 70746-3) and Halt protease and phosphatase inhibitor (Thermo Fisher Scientific, 78442), followed by centrifugation at 14,000 g for 10 min at 4°C. For nuclear co-IP, NE-PER™ nuclear and cytoplasmic extraction reagents (Thermo Fisher Scientific, 78835) supplemented with Halt protease and phosphatase inhibitor was used. After CER I/II lysis and centrifuge steps, nuclear pellet was washed with PBS once to remove remaining cytoplasmic fraction in supernatant before nuclear fraction lysis. Nuclear lysates were centrifuged at 14,000 g for 10 min at 4°C, followed by filtering with Amicon ultra-0.5 centrifugal filter unit (MilliporeSigma, UFC500396) to purify macromolecular components with molecule weight larger than 3 kDa. Lysate was then quantified by BCA protein assay. An equal amount of cleared lysate was mixed with Dynabeads Protein G (Thermo Fisher Scientific, 10004D) pre-conjugated with 1-3 μg of anti-ELMSAN1 (Novus Biologicals, NBP1-71912), anti-DLAT antibody (Thermo Fisher Scientific, MA5-24863), anti-PDHX (Proteintech, 10951-1-AP), anti-mouse IgG (Santa Cruz Biotechnology, sc-2025), anti-rabbit IgG (Novus Biologicals, NB810-56910). Equal amount (3 μg) of antibodies were used in the reciprocal nuclear co-IP experiment. The mix was incubated on a rotator at 4 °C overnight. The beads were then washed 4 times with lysis buffer and 2 times with PBS. For western blotting, beads were eluted with LDS sample buffer (Thermo Fisher Scientific, NP0008) and sample reducing agent (Thermo Fisher Scientific, NP0009), and boiled for 10 min. For the ELMSAN1 truncation co-IP experiment, truncations were constructed by Q5 site directed mutagenesis kit (NEB Biolabs, E0554S) on pLV-Puro-CMV>HA-hMIDEAS. For mass spectrometry, after wash steps, beads were frozen in -20°C. Cells (~80% confluency) were treated with indicated concentration of ISX9 in incubator for 2-4 hours, followed by lysis and overnight incubation steps supplemented with the same concentration of ISX9.

##### Recombinant protein purification and pull-down assay

Recombinant proteins GST-DLAT and 6xHis-ELMSAN1 were expressed by pET-GST/TEV/hDLAT and pET-6xHis/hELMSAN1 in BL21 (DE3) competent E. coli (NEB Biolabs, C2527) at 18°C for 18 hours after induction with 0.5 mM IPTG (MilliporeSigma, I6758). Cells were lysed by high-pressure homogenization using the Emulsiflex C3 system (Avestin). GST-DLAT protein was captured on glutathione sepharose 4B GST-tagged protein purification resin (Cytiva, 17075601) for pull-down assay. 6xHis-ELMSAN1 proteins were purified using Ni-NTA agarose (Qiagen, 30210), eluted with high concentration of imidazole buffer and further purified by filtering through a 50 kDa filter (MilliporeSigma, UFC905024). His-HA-GFP and His-HA-ELMSAN1 were *in vitro* translated by pT7CFE1-CHis-HA/GFP and pT7CFE1-CHis-HA/ELMSAN1 using 1-Step Human Coupled IVT Kit (Thermo Fisher Scientific, 88881). *In vitro* pull-down experiments were performed by incubating the mixture of resin-GST-DLAT with His-ELMSAN1 or His-HA-GFP and his-HA-ELMSAN1 in lysis buffer (Thermo Fisher Scientific, 87788) on rotator at 4 °C overnight. For the experiment with ISX9, different doses of ISX9 were added to the mixture for incubation overnight. The resins were then washed 4 times with lysis buffer and eluted for western blot.

**Immunoblotting (western blotting, WB)**

For whole cell lysate, cells were lysed in ice-cold 1x RIPA buffer (MilliporeSigma, R0278) with Halt protease and phosphatase inhibitor. After being cleared by centrifugation at 14,000 g for 15 min at 4°C, cleared lysate was quantified by BCA protein assay. An equal amount of cleared lysate was mixed with LDS sample buffer and sample reducing and boiled for 10 min. Co-IP samples for WB were prepared as described above. Proteins were loaded in gels (Bio-Rad, 4568095) and transferred onto PVDF membranes (Thermo Fisher Scientific, LC2005). Membranes were blocked in TBS (LI-COR, 927-60001) and incubated with anti-GAPDH (CST, 2118S, 1:1000), anti-H3 (Abcam, ab1791, 1:5000), anti-H3K27ac (Abcam, ab4729, 1:1000), anti-DLAT (CST, 12362S, 1:1000), anti-PDHX (Proteintech, 10951-1-AP, 1:1000), anti-ELMSAN1 (Novus Biologicals, NBP1-71912), anti-HA (CST, 3724, 1:1000), anti-Lamin A/C (Millipore Sigma, MABT538, 1:1000), anti-COX IV (Proteintech, 11242-1-AP, 1:1000), anti-HDAC1 (CST, 5356, 1:1000), anti-HDAC2 (CST, 5113, 1:1000), anti-H2A (CST, 12349, 1:1000), anti-H2AZ (CST, 2718, 1:1000) at 4°C overnight, followed by staining secondary antibodies (LI-COR, 926-32211, 926-68020). For staining PDHX in co-IP experiments, anti-rabbit IgG (Light-Chain Specific) (HRP Conjugate) (CST, 93702S, 1:1000) was used as secondary antibody. Signals were detected and quantified using the LI-COR Odyssey CLx imaging system or film.

**Mass spectrometry proteomics**

Sample Preparation. The samples were prepared for proteomic analysis using established protocols at Harvard Center for Mass Spectrometry. IP samples were received on magnetic beads and any remaining buffer solution was removed. Samples were first gently washed 3 times with 100 µL of fresh PBS, and then were redissolved in 100 µL of 50 mM TEAB and incubated at 95ºC for 5 min. After samples returned to room temperature, they were trypsin digested for 3 hours at

37°C. Digested samples were cleaned up with C18 Spin Tips (Pierce, Thermo Fisher Scientific) and reconstituted with 0.1% formic acid prior to LC-MS/MS analysis.

LC-MS/MS Method. Samples were analyzed by Q Exactive HF-X High Resolution Orbitrap (Thermo Fisher Scientific) coupled with Ultimate 3000 nanoLC (Thermo Fisher Scientific).

Analysis of Proteomics Data. Raw data was then submitted for analysis in Proteome Discoverer 3.0 software (Thermo Fisher Scientific). The MS/MS Data was searched against the UniProt reviewed Homo sapiens database as well as other known contaminants such as human keratins and common lab contaminants. At least one unique peptide per protein group is required for identifying proteins.

##### Cell-based proteome integral solubility alteration

Cell preparation. PDXC-OS052 cells grown to ~75% confluency were trypsinized, washed once with PBS, and suspended in fresh media. ISX9 (50 µM), MGCD0103 (5 µM), I+M, or DMSO were incubated with cell suspension at 37°C for 30 min. An equal volume of each sample was aliquoted across ten PCR tubes and heated to a different temperature from 48°C-58°C for 3 min in an Eppendorf Mastercycler Pro S. The samples were allowed to cool to room temperature for 5 min. An equal volume from each PCR tube was pooled in a fresh tube. The pooled sample was spun at 300 g for 3 min to pellet the cells. The cells were washed once with ice-cold PBS and suspended in an extraction buffer (PBS with 1.5 mM $MgCl_2$, 0.5% NP-40, 1X protease inhibitor [Peirce protease inhibitor mini tablets]). The samples were incubated for 10 min at 4°C and spun at 21,000 g for 90 min. ~15 µg of soluble protein were collected and prepared for LC-MS/MS analysis.

##### PDC enzyme activity and protein quantity assay

An equal number of cells were used in each experiment. > 5 million cells were usually needed to generate a visible result from the nuclear fraction. For ISX9 treatment, 143B cells grown in 150 mm dishes (~80% confluency) were treated with indicated concentration of ISX9 and the same volume of DMSO in incubator. Non-nuclear and nuclear fractions were isolated using NE-PER™ nuclear and cytoplasmic extraction reagents with the same concentration of ISX9. Then, each fraction was filtered by Amicon ultra-0.5 centrifugal filter unit to purify macromolecular components. Mitochondrial proteins were isolated using the mitochondria Isolation Kit (Abcam, ab110170). Pyruvate dehydrogenase enzyme activity dipstick assay kit (Abcam, ab109882) and pyruvate dehydrogenase protein quantity dipstick assay kit (Abcam, ab109883) were used to measure PDC enzyme activity and protein quantity. Briefly, for activity assay, mitochondrial extracts or concentrated nuclear extracts were loaded in a well of 96-well plate and PDC can be immunocaptured in its native form with an anti-PDC-E2 (DLAT) antibody immobilized on a defined section of a dipstick. The PDC activity is measured and visualized by coupling PDC-dependent production of NADH to the reduction of NBT in the presence of excess diaphorase, forming an insoluble intensely colored precipitate at the capture line. 5%-20% non-nuclear fractions was loaded and developed for 10-20 min. ~50% nuclear fractions were loaded and developed for ~1 hour. The developing time between samples was the same. For quantity assay, mitochondrial extracts or concentrated extracts were loaded in a well of 96-well plate and PDC can be immunocaptured in its native form with anti-DLAT/PDHX antibodies immobilized on a defined section of a dipstick for visualization. Quantification was performed by ImageJ.

**Nuclear acetyl-CoA metabolite extraction**

An equal number of cells were used in each experiment. 5-10 million cells were usually needed to generate a reliable acetyl-CoA result from the nuclei. 143B cells grown in 150 mm dishes (60~80% confluency) were treated with 100-300 µM ISX9 and the same volume of DMSO in an incubator followed by nuclear isolation by NE-PER™ nuclear and cytoplasmic extraction reagents. Briefly, dishes were placed on ice and the medium was removed completely. After washing with PBS and removing any residual liquids from the dish completely, hypotonic buffer (CER I) was added directly into the dish. Cells were scraped, pipetted up and down several times, and transferred to a tube on ice. CER II was then added to break the cell membrane for 1 min. Nuclei were pelleted by centrifuging at 3,000 g for 5 min. Pellets were washed with CER I and centrifuged at 3,000 g for 5 min. After removing the supernatant, the residual buffer was removed by 10 µl pipetting. Purified nuclei were resuspended in -80°C pre-cold 800-1000 µl of 80:20 UHPLC-grade methanol (MilliporeSigma, 900688) and UHPLC-grade $H_2O$ (MilliporeSigma, 900682), followed by repeatedly pipetting and vigorous vortex. Samples were then placed in liquid nitrogen for 10 min and thawed on ice for 10 min, followed by pipetting and vigorous vortex. The freeze-thaw cycle was repeated twice. Samples were centrifuged at 14,000 g to remove cell debris, lipids, and proteins. The supernatant was transferred to new tubes and kept at -80°C before LC-MS analysis.

For the analysis with SILEC-SF internal standard, we generated 143B SILEC cells as reported[50,64]. Briefly, 143B cells were cultured for > 10 passages in DMEM without pantothenate

(United States Biological, D9800-02B) supplemented with $^{15}N_1{}^{13}C_3$-vitamin B5 (MilliporeSigma, 705837) and charcoal/dextran stripped FBS (Gemini Bioproducts, 100-119-500). FBS concentration was decreased from 10% to 3% and $^{15}N_1{}^{13}C_3$-vitamin B5 concentration was increased from 1 µg/ml to 3 µg/ml in late passages to increase the efficiency of isotope labeling on acetyl-CoA. The labeling efficiency of nuclear $^{15}N_1{}^{13}C_3$ acetyl-CoA was confirmed to be 99.2%. An equal number of SILEC 143B cells were added to each sample of DMSO and ISX9 treated 143B cells (without isotope label) prior to nuclei isolation for metabolite extraction.

**Mass spectrometry analysis of acetyl-CoA**

Samples were analyzed by LC-MS on a Vanquish LC coupled to an ID-X MS (Thermo Fisher Scientific). Glutamate or SILEC-SF were used as the internal standard. 5 µL of sample or standard was injected on a ZIC-pHILIC peek-coated column (150 mm x 2.1 mm, 5-micron particles, maintained at 40°C, MilliporeSigma). Buffer A was 20 mM Ammonium Carbonate, 0.1% Ammonium hydroxide in water and Buffer B was Acetonitrile 97% in water. The LC program was as follows: starting at 93% B, to 40% B in 19 min, then to 0% B in 9 min, maintained at 0% B for 5 min, then back to 93% B in 3 min, and re-equilibrated at 93% B for 9 min. The flow rate was maintained at 0.15 mL/min, except for the first 30 s where the flow rate was uniformly ramped from 0.05 to 0.15 mL/min. Data was acquired on the ID-X in negative mode at 120,000 resolution, an RF lens at 30%, and a m/z range of 65 to 1000. To improve the acetyl-CoA signal, data was also acquired in tSIM mode on the $[M-2H]^{2-}$ ion for acetyl-CoA, with an isolation of 4 m/z. Extracted ion chromatogram (within 5ppm) for the acetyl-CoA ion in the tSIM signal was integrated using Tracefinder (TF 4.1, Thermo Fisher Scientific).

**Generation of CRISPRa cell line**

143B cells were employed to generate CRISPRa cell lines. Cells were first transduced with a lentiviral vector expressing pHRdSV40-dCas9-10xGCN4_v4-P2A-BFP (Addgene, #60903) and purified for $BFP^+$ cells by flow cytometry. Next, cells were transduced with pHRdSV40-scFv-GCN4-sfGFP-VP64-GB1-NLS (containing SunTag-binding single-chain antibody fused with VP64) (Addgene, #60904) and purified for $BFP^+GFP^+$ cells by several rounds of flow cytometry sorting to achieve >99.9%. The activity of CRISPRa line was validated by transducing with positive control sgRNAs followed by qPCR analysis of gene expression: *SLC4A1* and *POU5F1*. sgRNA targeting *Gal4* (sg*Gal4*) was used as a control.

#### Amplification of CRISPRa pooled libraries

Human genome-wide CRISPRa-v2 libraries (Addgene, #83978) were obtained from Addgene. ~200 ng pooled libraries were transformed into MegaX DH10B cells (Thermo Fisher Scientific, C640003) by electroporation (setting: 2.0 kV, 200 ohms, 25 µF in 0.1cm cuvette) to achieve a transformation efficiency of >1,000 colonies/sgRNA. Colonies were grown on LB agar plates overnight and harvested on the next day, followed by plasmid extraction using the NucleoBond Xtra Maxi kit (Macherey-Nagel, 740424). Next, the sgRNA target regions from the amplified libraries were PCR amplified using primers with Illumina adaptors and sequenced by next-generation sequencing to determine the sgRNA distribution of libraries.

#### Genome-wide CRISPRa screen

The CRISPRa-v2 libraries were transduced in duplicate into 143B CRISPRa cells in T175 flasks at the MOI ~0.3 (virus volume was determined by conducting a small-scale pilot test). 2 days after transduction, cells were selected by 5 µg/ml puromycin for 2 days, followed by 2 days culture without puromycin. Then, cells were trypsinized and split into 3 groups for treatment. $1x10^8$ cells were seeded for each group in T500 flasks, which allowed a library coverage of more than 1,000 cells per sgRNA. Cells were treated with DMSO and ISX9 (3 days of 30 µM, 1 day of 20 µM and 2 days of 5 µM) for 6 days before harvesting. The concentration of ISX9 was optimized and adjusted to keep cells in a slow-growing or non-proliferative status, resulting in 6-8 fewer doublings than DMSO-treated cells. Finally, $6x10^7$~$1x10^8$ cells from each group of each duplicate were harvested. Genomic DNA was isolated using NucleoSpin Blood Maxi kit (Macherey-Nagel, 740950). Next, sgRNA target regions were PCR amplified with Illumina adaptors and indexes using NEBNext® Ultra™ II Q5® Master Mix (NEB Biolabs, M0544X) and products were purified with DNA Clean & Concentrator-100 (Zymo Research, D4030). sgRNA target regions (260-280bp) were further enriched by gel extraction (Qiagen, 28704) after running agarose gel. Purified PCR products were measured by Qubit and analyzed by TapeStation before sending for next-generation sequencing on Illumina NextSeq 2000 with 50-bp single-end-read settings.

#### CRISPRa screen analysis

The CRISPR screening data were processed and analyzed using the MAGeCK algorithms[65]. Raw sequencing data were pre-processed using MAGeCK to obtain the read counts for each sgRNA, then the MAGeCK robust rank aggregation (RRA) method was used to identify negative and positive selected genes. Briefly, MAGeCK constructed a linear model to compute the variance of guide RNA (gRNA) read counts and assessed the difference in gRNA abundance between the control and treatment conditions. The selection of genes was evaluated from the rankings of gRNAs (by their *P* values) using the α-RRA algorithm. For each gene, α-RRA assigned *P* values for both positive and negative selection.

**RNA-seq and analysis**

Total RNA was isolated using Direct-zol RNA MiniPrep Kit (Zymo Research, R2052), and mRNA was purified using the NEBNext® Poly(A) mRNA Magnetic Isolation Module (New England Biolabs, E7490). Libraries were generated using the NEBNext® Ultra™ RNA Library Prep Kit for Illumina® (New England Biolabs, E7530). Samples were sequenced on Illumina NextSeq 2000 with 50-bp single-end-read or 150-bp paired-end-read settings.

RNA-seq reads were mapped to the hg19 reference genome using STAR v. 2.7.3a[66]. Read counts for individual genes were produced using the count feature in HTSeq v. 0.6.0[67]. Differential gene expression analysis was performed using the edgeR package v. 3.36.0[68] after normalizing read counts and including only genes with count per million reads (CPM) > 1 for one or more samples. Differentially expressed genes were defined based on the criteria of fold change (FC) > 2 in expression value and false discovery rate (FDR) < 0.05. Heatmaps and PCA plots were created using R packages ComplexHeatmap v. 2.10.0[69] and ggplot2 v. 3.3.6, respectively. Analysis of gene set enrichment was performed using Enrichr[70] and GSEA[71].

**ATAC-seq**

PDXC-OS052 and Rh30 cells were treated for 6 days. Then, cells were trypsinized, and approximately 50,000 cells were used to perform the ATAC assay as previously described[72] with duplicates for each sample. Briefly, cells were first lysed in buffer (10 mM Tris-HCl pH7.4, 10 mM NaCl, 3 mM $MgCl_2$, and 0.1% Igepal CA-630). Then, cell pellets were incubated with transposition reaction mix (25 µl Tagment DNA Buffer, 2.5 µl TDE1 Tagment DNA Enzyme, and 22.5µl nuclease-free H2O) (Illumina, 20034197) for 30 min at 37°C in a thermomixer at 300 rpm, followed by PCR reaction for 18 cycles. All samples were sequenced on Illumina NextSeq 2000 with 50-bp single-end-read settings.

#### CUT&Tag

PDXC-OS052 cells were treated with DMSO and 100 µM ISX9 for 3 hours for CUT&Tag of H3K27ac using the CUT&Tag-IT™ Assay Kit (Active Motif, 53160) with H3K27ac antibody (Abcam, ab4729). Briefly, cells were first harvested by Accutase and incubated with concanavalin A beads, and then incubated with primary antibodies followed by secondary antibody. The chromatin is digested by tagmentation using the protein A-Tn5 (pA-Tn5) transposase. After DNA purification and PCR amplification with adaptors, samples were sequenced on Illumina NextSeq 2000 with 50-bp single-end-read settings.

#### ATAC-seq and CUT&Tag analyses

Sequencing reads were aligned to the hg19 reference genome using BWA v. 0.7.17-r1188[73] with command bwa sampe. Peaks were called using HOMER v. 4.10.3[74] with default parameters. DiffBind v. 3.4.11 R package[75] was used for differential peak analysis based on the cutoffs of FC > 2 and FDR < 0.05. Transcription factor binding sequence motif enrichment was analyzed using HOMER (v. 4.11) motif analysis[74]. Differential ATAC and H3K27ac peaks were defined by FC > 2 and FDR < 0.05. Genomic track was visualized by Integrative Genomics Viewer (v. 2.11.9)[76].

#### Secondary transplantation

PDX tumors were harvested from primary transplanted mice after treatment with vehicle or I+M. Tumor cells were isolated as described above and stained with anti-human HLA-A,B,C (BioLegend, clone W6/32, 1:100). Flow cytometry determined the percentage of human cells (human HLA-A,B,C positive cells). Human cell percentage generally did not differ between vehicle and I+M treated tumors. An equal number of live human cells (1-3x$10^6$) from each group were transplanted subcutaneously into secondary recipient NSG mice. Mice were monitored every week, and tumor volumes were measured as above. Mice were euthanized according to the same standards shown above.

#### Limiting dilution analysis in secondary transplantation

A serial number of tumor cells from primarily treated mice were transplanted into secondary recipient NSG mice. The number of palpable/visible tumors was counted 5-6 weeks after transplantation. TIC frequency and significance were calculated by ELDA software[77].

**Toxicity study *in vivo***

Four-week-old C57BL/6J (The Jackson Laboratory, 000664) mice were administered six days per week with indicated small molecules for three weeks. Body weights were monitored every week. Peripheral blood was retro-orbitally collected every week and complete blood was counted using Element HT5 Auto Hematology analyzer (Heska). Mice were euthanized at the endpoint (3 weeks). Hearts, lungs, spleens, kidneys, and livers were weighed. The analyses and colony formation assay of mesenchymal stromal cells were performed as stated below.

**Flow cytometry and cell sorting**

For isolation of mouse mesenchymal stromal cells in the toxicity studies, bone fractions were cut and digested with 0.25% collagenase I (Stem Cell Technologies, 07902), 2 mg/ml dispase (Thermo Fisher Scientific, 17105041) and 12 U/ml DNase I (Thermo Fisher Scientific, 90083). Red blood cells were lysed in ACK buffer. Cells were stained with anti-CD45 (BioLegend, clone 30-F11, 1:100), anti-CD31 (BD Biosciences, clone MEC 13.3, 1:100), anti-Ter119 (BD Biosciences, clone TER-119, 1:100), anti-Sca-1 (BioLegend, clone D7, 1:100), anti-CD51 (BD Biosciences, clone RMV-7, 1:100), and anti-CD140a (Thermo Fisher Scientific, clone APA5, 1:100). DAPI was used for viability staining. Cells were sorted by BD FACSAria flow cytometer. For analyzing and sorting tumor samples, cells were isolated as described above and stained with anti-human HLA-A,B,C for separation of human tumor cells, anti-CD44 (BioLegend, clone BJ18, 1:100) and anti-CD133 (BD Biosciences, clone W6B3C1, 1:100) for analyzing TIC. DAPI was used for viability staining. Cells were analyzed by BD LSR II flow cytometer.

**Colony formation unit assay**

3,000 $CD31^{-}CD45^{-}Ter119^{-}$ mesenchymal stromal cells sorted from bone marrow were seeded in a 6-well plate in MEMα supplemented with 20% FBS, 1% PS, and 1% Glutamine. Colonies were visualized and counted 26 days after culture by staining crystal violet.

#### Correlation/dependency analyses

Between neuronal and proliferation signature in 10 sarcoma lines. Average RNA-seq expressions (RPKM) of neuronal genes *NEUROD1*, *MAP2*, *NEFM*, *SYN1*, *SYP*, *TUBB3* and proliferation genes *MKI67*, *AURKA*, *CCNA2*, *CDK1*, *FOXM1* were used to calculate Z-scores and scatterplot.

Between ELMSAN1 expression and drug sensitivity. RNA expression of *ELMSAN1* and ISX9, MGCD0103 drug sensitivity data were downloaded from the DepMap database. GraphPad Prism 9 was used to generate scatterplots and calculate simple linear regression and correlation in corresponding cancer lineages using all cell line data in that lineage.

Chronos dependency score of cell-of-origin TF. Chronos scores of all cell lines [CRISPR (Public 23Q2+Score, Chronos), *n* = 1,096] were downloaded from the DepMap database. Chronos score indicates the dependency of a cell line on a gene. A gene score of 0 means not essential while a score of -1 corresponds to the median of all common essential genes. GraphPad Prism 9 was used to generate plots. The enriched cancer lineage was colored, and the corresponding TF and *P* value (from the DepMap database) were indicated.

#### Patient survival analysis using *ELMSAN1* cancer signature

*ELMSAN1* cancer signature (the expression of *IER3*, *MMP14*, *GJA1*, *COL6A2*, *COL9A3*, *CTHRC1*, *KRT8*, *ITGB3*, *LRP10*, *FBLIM1*, *GSN*, *MRC2*, *RTN1*, *H1F0*) was created by overlapping top upregulated DEGs in *ELMSAN1*-OE and top downregulated DEGs in *ELMSAN1*-KD. Patient survival data and the RNA-seq data were downloaded from the TCGA database. To test if the *ELMSAN1* cancer signature predicts survival, we first computed the average expression of the signature in each tumor based on the normalized RNA-seq data. Next, we stratified the patients into two groups according to the average expression of the signature: high or low expression corresponded to the top or bottom 25% of the population. We used a log-rank test to examine if there was a significant difference between patient groups in terms of their survival. The survminer R package was used to draw Kaplan Meier survival plot. Multi-variate analysis between the *ELMSAN1* cancer signature and proliferation score and EMT score was analyzed using the TCGA database.

#### Protein and small molecule docking

Molecular docking of DLAT protein (PDB ID: 6H55) and ELMSAN1 (PDB ID: 6Z2J_3) was conducted using the ZDock sever[78]. Next, LeDock[79,80] was employed to predict the interaction between the docked protein complex and ISX9 based on the identified binding pockets by POCASA[81].

#### QUANTIFICATION AND STATISTICAL ANALYSIS

Statistical and graphical data analyses were performed using GraphPad Prism 9 or other indicated software. The number of samples, details of statistical test used are provided in figures or figure legends.

#### Supplemental Figure 1

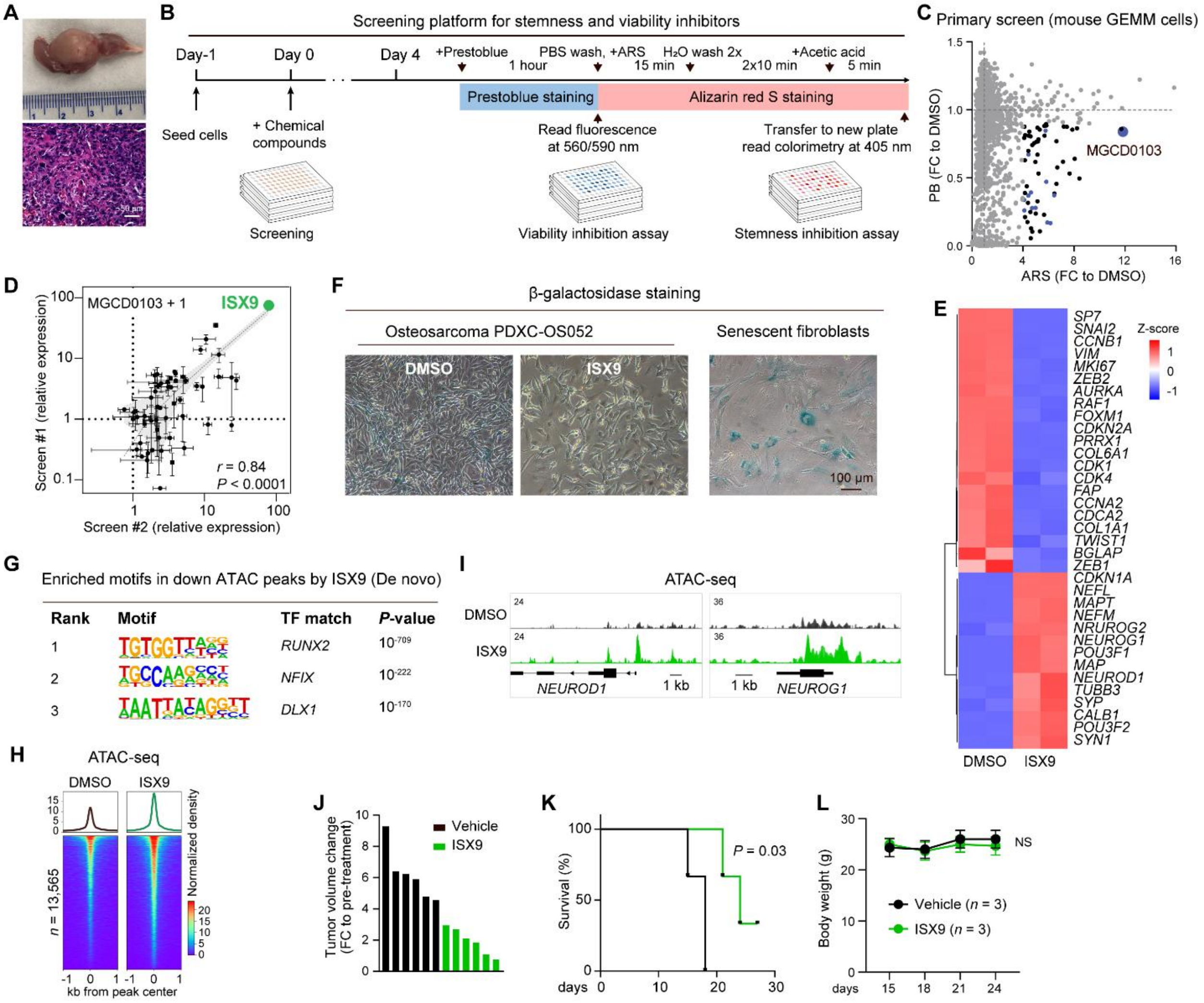


**Figure S1. Therapeutic cancer cell reprogramming by small molecules, related to Figure 1.**

(A) Osteosarcoma GEMM cells OS2674 formed tumors after subcutaneously transplanting into immunocompromised mice (upper: tumor formation around femur, bottom: H&E staining of the tumor). Scale bar, 50 µm.

(B) Procedures depicting the screening platform for stemness and viability inhibitors.

(C) Effect of each small molecule on ARS and cell viability in the primary screen. HDACi are highlighted in blue.

(D) qPCR analysis of small molecule effect on *IBSP* expression based on MGCD0103 in 143B cells. Duplicate screens exhibited a strong correlation. $n = 2$.

(E) Heatmap of representative MNLT gene expression under DMSO and ISX9 treatments for 6 days with unsupervised clustering with normalized expression as Z-score. $n = 2$.

(F) β-galactosidase staining of PDXC-OS052 under DMSO and ISX9 treatments for 6 days. Senescent fibroblasts were used as a positive control. Scale bar, 100 µm.

(G) De novo analysis of the TF motif enrichment of downregulated ATAC peaks by ISX9.

(H) Heatmaps and average profiles of ATAC-seq peaks increased by ISX9. $n$ = 2.

(I) Genomic tracks of representative increased ATAC-seq peaks. $n$ = 2 tracks overlaid.

(J-L) Waterfall plot of individual tumors ($n$ = 6, J), mouse survival ($n$ = 3, K), and mouse body weights ($n$ = 3, L) under vehicle and ISX9 (50 mg/kg, twice a day) treatment.

Data are presented as mean ± SEM. Statistical significance was determined using simple linear regression (D), log-rank (Mantel-Cox) test for survival curves (K), or two-tailed unpaired t-tests (L).

#### Supplemental Figure 2

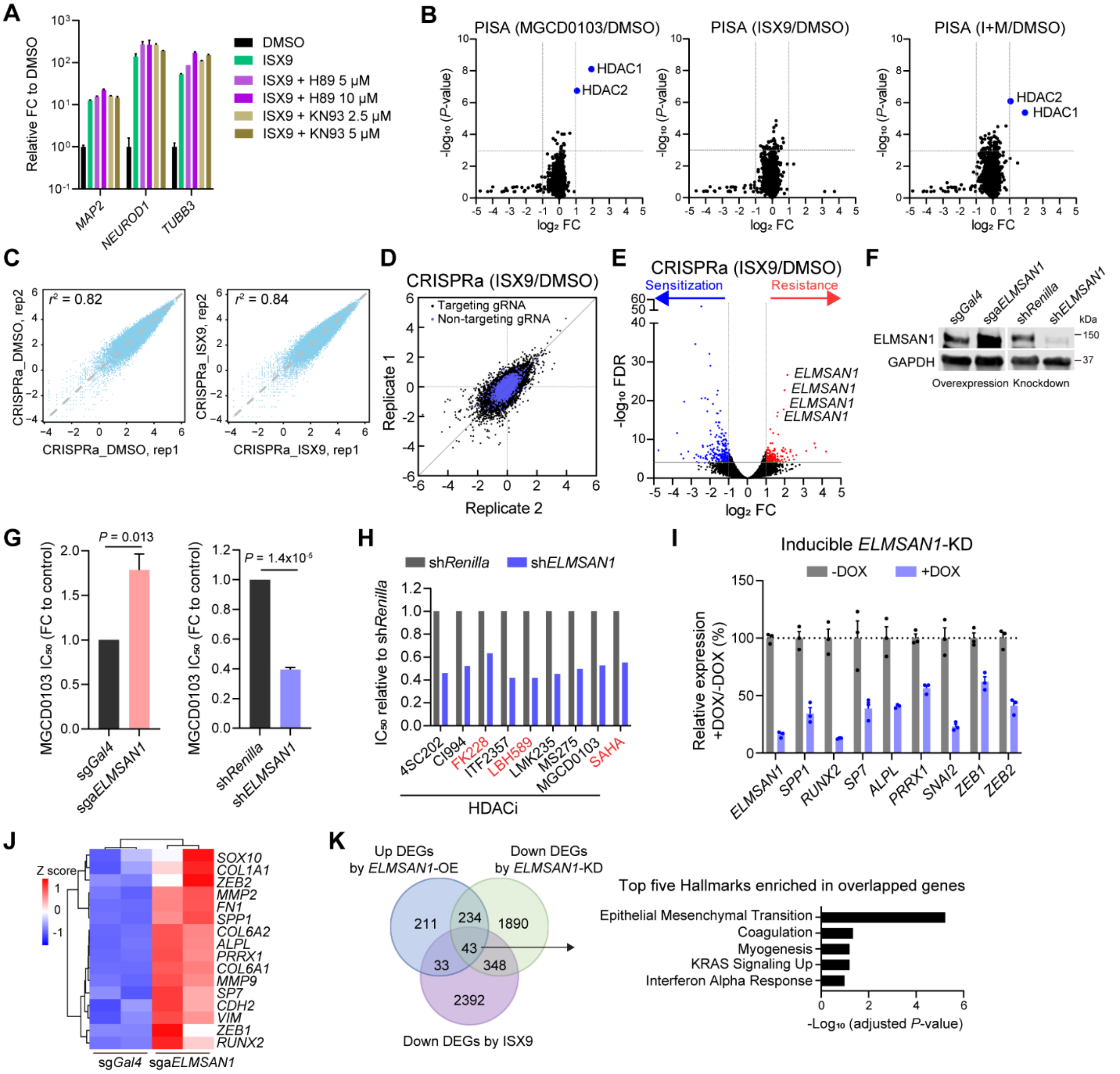


**Figure S2. Genome-wide CRISPR activation screen identifies ELMSAN1 as the reprogramming mediator, related to Figure 2.**

(A) qPCR analysis of neuronal gene expression under indicated treatment. H89 is a protein kinase A inhibitor; KN93 is a CaMKII inhibitor. *n* = 2.

(B) Volcano plots of the PISA assay show the differential thermal stability of proteins affected by ISX9, MGCD0103, and I+M in PDXC-OS052. Despite further confirming HDAC1 and HDAC2 as MGCD0103 targets, PISA could not reveal a faithful target for ISX9. *n* = 4.

(C) Scatterplots showing the correlation ($r^2$) between duplicate samples in the CRISPRa screen.

(D) Representative scatterplot depicting the correlation between duplicate samples comparing ISX9 with DMSO. Blue dots indicate non-tarting sgRNA, which are clustered around 0.

(E) Volcano plot showing the individual sgRNA in CRISPRa screen comparing ISX9 vs DMSO. Lines demarcate threshold: FC = 2 and FDR = 0.0001. Blue dots represent genes that synergized treatment, whereas red dots represent genes that antagonized treatment. *n* = 2.

(F) WB of ELMSAN1 expression upon KD and OE.

(G) $IC_{50}$ of MGCD0103 upon KD and OE of *ELMSAN1* relative to control. *n* = 3, except ELMSAN1-KD (*n* = 2).

(H) $IC_{50}$ of HDACi upon *ELMSAN1*-KD. Red highlights three FDA-approved HDACi for cancer treatment.

(I) qPCR analysis of mesenchymal/osteogenic gene expression upon *ELMSAN1*-KD. *n* = 3.

(J) RNA-seq gene expression heatmap of mesenchymal/osteogenic genes upon *ELMSAN1*-OE. *n* = 2.

(K) Left: Venn diagram of upregulated DEGs in ELMSAN1-OE, downregulated DEGs in ELMSAN1-KD, and downregulated DEGs by ISX9. Right: enrichment analysis of the overlapped 43 genes.

Data are presented as mean ± SEM. Statistical significance was determined using two-tailed unpaired t-tests (B, E, G), simple linear regression (C), or Benjamini-Hochberg method (K).

#### Supplemental Figure 3

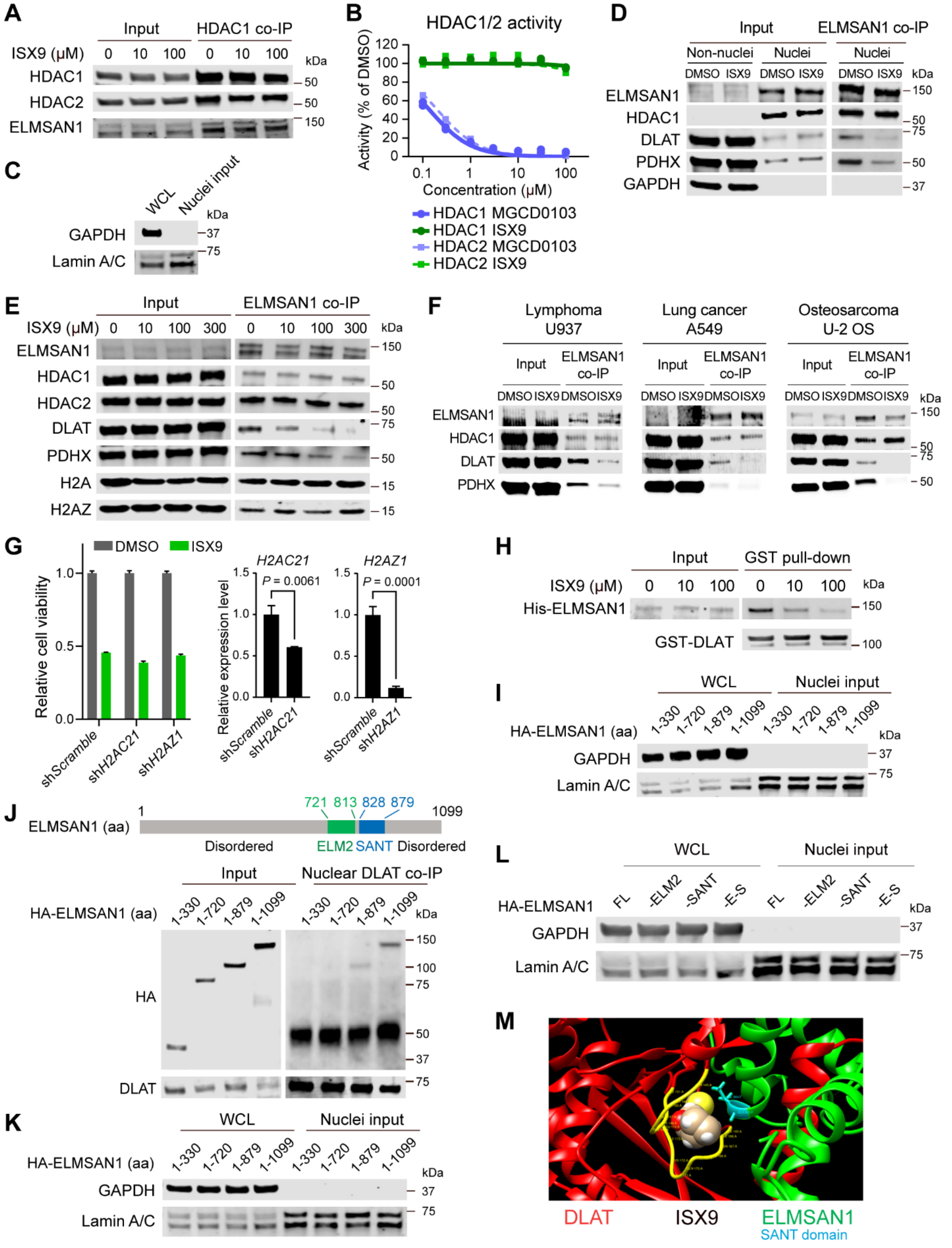

**Figure S3. ELMSAN1 binds to PDC which can be pharmacologically disrupted, related to Figure 3.**

(A) HDAC1 co-IP assay confirmed unchanged HDAC1-ELMSAN1 binding upon ISX9 treatment.

(B) HDAC1/2 enzyme activity assay assessing the MGCD0103 and ISX9 effect on inhibiting HDAC1/2 activity. $n$ = 2.

(C) Western blot confirmed the isolation of nuclei sample in Figure 3B. GAPDH is a cytoplasm marker. Lamin A/C is a nucleus marker.

(D) co-IP validation of PPI inhibition in the nucleus. 143B cells were treated with ISX9 for 3 hours before co-IP. Input suggests ELMSAN1 in the nucleus and the majority of DLAT and PDHX in the non-nuclear fractions.(E) ELMSAN1 co-IP confirmed the unchanged HDAC1/2 binding, lost DLAT/PDHX binding, and mildly enhanced H2A, H2AZ binding upon ISX9 treatment.

(F) ELMSAN1 co-IP showed disrupted ELMSAN1-DLAT/PDHX interaction by ISX9 treatment for 3 hours in lymphoma, lung cancer and osteosarcoma cells. ELMSAN1-HDAC1 interaction is not changed.

(G) *H2AC21* and *H2AZ1* knockdowns did not change ISX9's effect on reducing cell growth. Knockdown efficiency was shown on the right. $n$ = 3 except for H2AC21 sh*Scramble* $n$ = 2.

(H) Pull-down of recombinant GST-DLAT proteins with recombinant His-ELMSAN1 protein in the presence of ISX9 at different doses *in vitro*.

(I) Western blot confirmed the isolation of nuclei sample in Figure 3E.(J) Nuclear DLAT co-IP with HA-ELMSAN1 truncations expressed in 143B cells to identify critical PPI domains. 721-813aa is the ELM2 domain, and 828-879aa is the SANT domain in the ELMSAN1 protein.

(K) Western blot confirmed the isolation of nuclei sample in Figure S3J.

(L) Western blot confirmed the isolation of nuclei sample in Figure 3F.

(M) Docking analysis of ISX9 interaction on ELMSAN1-DLAT. ELMSAN1 is green with the interaction region highlighted as cyan, which is within the SANT domain. DLAT is red with the interaction region highlighted as yellow. ISX9 was predicted to dock between the cyan-colored SANT domain and the yellow-colored DLAT.

#### Supplemental Figure 4

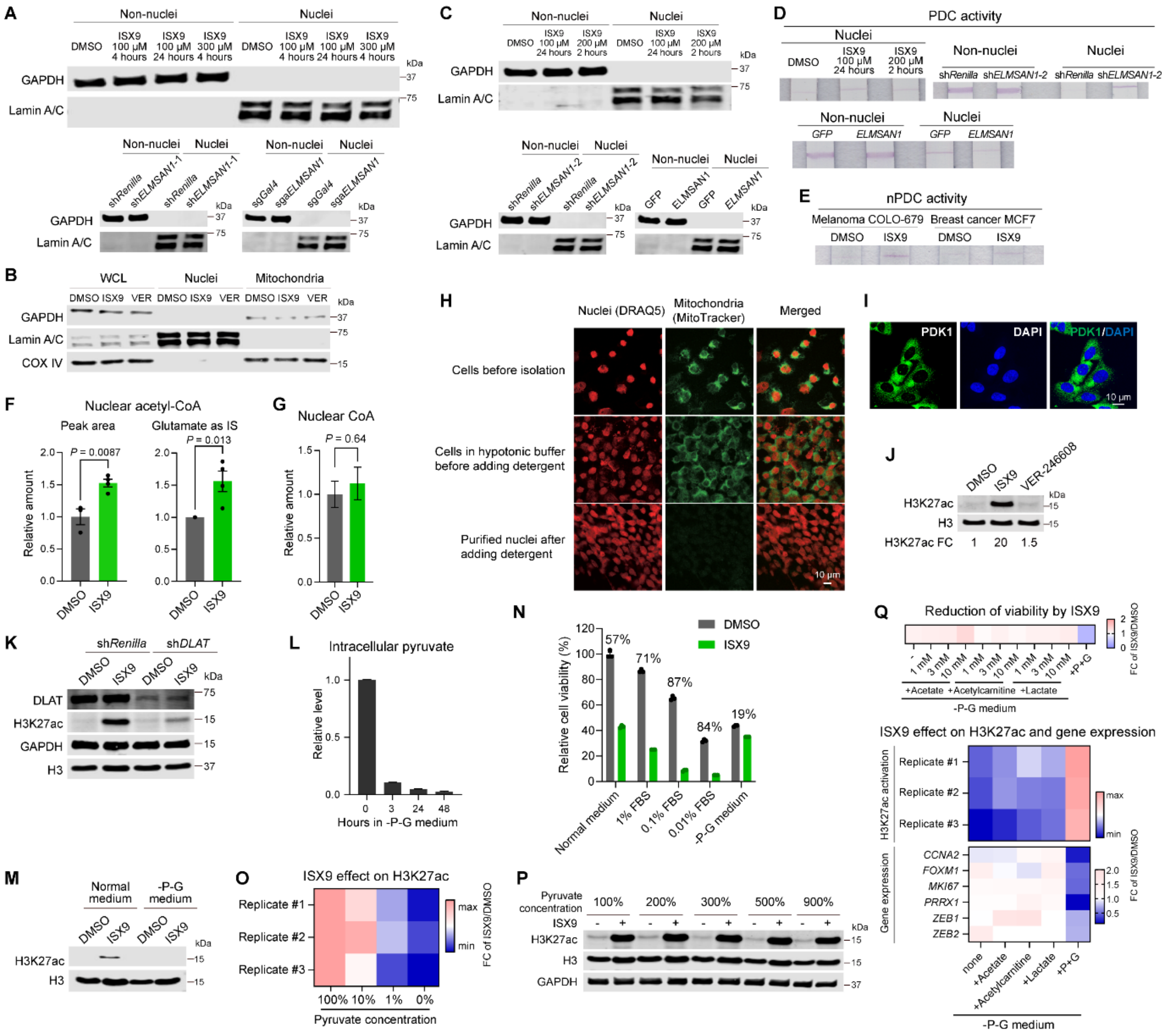

**Figure S4. Inhibiting ELMSAN1-PDC interaction de-represses nucleus-restricted PDC, related to Figure 4.**

(A) Western blot confirmed the isolation of nuclei in the PDC assay in Figure 4A.

(B) Western blot confirmed the isolation of nuclei and mitochondria in the PDC assay in Figure 4B. COX IV is a mitochondria marker.

(C) Western blot confirmed the isolation of nuclei in the PDC assay in Figure S4D.

(D) PDC activity under indicated conditions with the same cell number. The non-nuclear fraction (including cytosol and mitochondria) shows no difference in PDC activity between groups.

(E) Dipsticks of nuclear PDC activity under DMSO and ISX9 treatment for 3 hours in melanoma and breast cancer cells.

(F) LC-MS analysis of nuclear acetyl-CoA levels after DMSO and ISX9 treatment for 3 hours in 143B cells by measuring peak area ($n \geq 3$) or using glutamate as internal standard ($n = 4$ independent experiments).
(G) LC-MS analysis of nuclear CoA levels after DMSO and ISX9 treatment ($n \geq 4$).
(H) Immunofluorescence images of co-staining mitochondria (MitoTracker) and nuclei (DRAQ5) during the nuclear isolation process. Scale bar, 10 μm.
(I) Immunofluorescence images in 143B cells showing low or absent presence of PDK1 in the nucleus. Scale bar, 10 μm.
(J) WB of H3K27ac under DMSO, ISX9, and VER-246608 treatments for 3 hours.
(K) WB of indicated proteins under DMSO and ISX9 treatment for 3 hours in control and *DLAT*-KD cells. Quantification is shown in Figure 4H.
(L) Relative intracellular pyruvate concentration measured by pyruvate-Glo after culturing cells in -P-G medium for 0, 3, 24, and 48 hours. $n = 3$.
(M) WB of H3K27ac under DMSO and ISX9 treatments for 3 hours in normal medium and pyruvate-limiting medium.
(N) Relative cellular viability under DMSO and ISX9 treatment in different mediums, including normal medium, -P-G medium, and medium with different FBS concentrations. The percentage of cell viability reduction by ISX9 was indicated. -P-G medium and low FBS (0.01%-0.1%) medium reduced baseline cell growth to a similar level, but they changed ISX9 effect oppositely. -P-G medium abrogated ISX9 effect whereas the low FBS medium sensitized ISX9 effect, indicating that the abrogated ISX9 effect in -P-G medium was not attributed to slower baseline cell growth. $n = 3$.
(O) The activation of H3K27ac by ISX9 showed a pyruvate/glucose concentration-dependent manner when pyruvate was not saturated. The results of three replicates are shown. 100% refers to normal concentration, and 0% refers to -P-G medium.
(P) The activation of H3K27ac by ISX9 showed a pyruvate/glucose concentration-independent manner when pyruvate was saturated.
(Q) The effect of ISX9 on reducing viability, activating H3K27ac and changing MNLT gene expression under -P-G medium supplemented with acetate, acetylcarnitine, and lactate. $n = 3$.
Data are presented as mean ± SEM (F, G, L, N) or mean (O, Q). Statistical significance was determined using two-tailed unpaired t-tests (F, G).

### Supplemental Figure 5

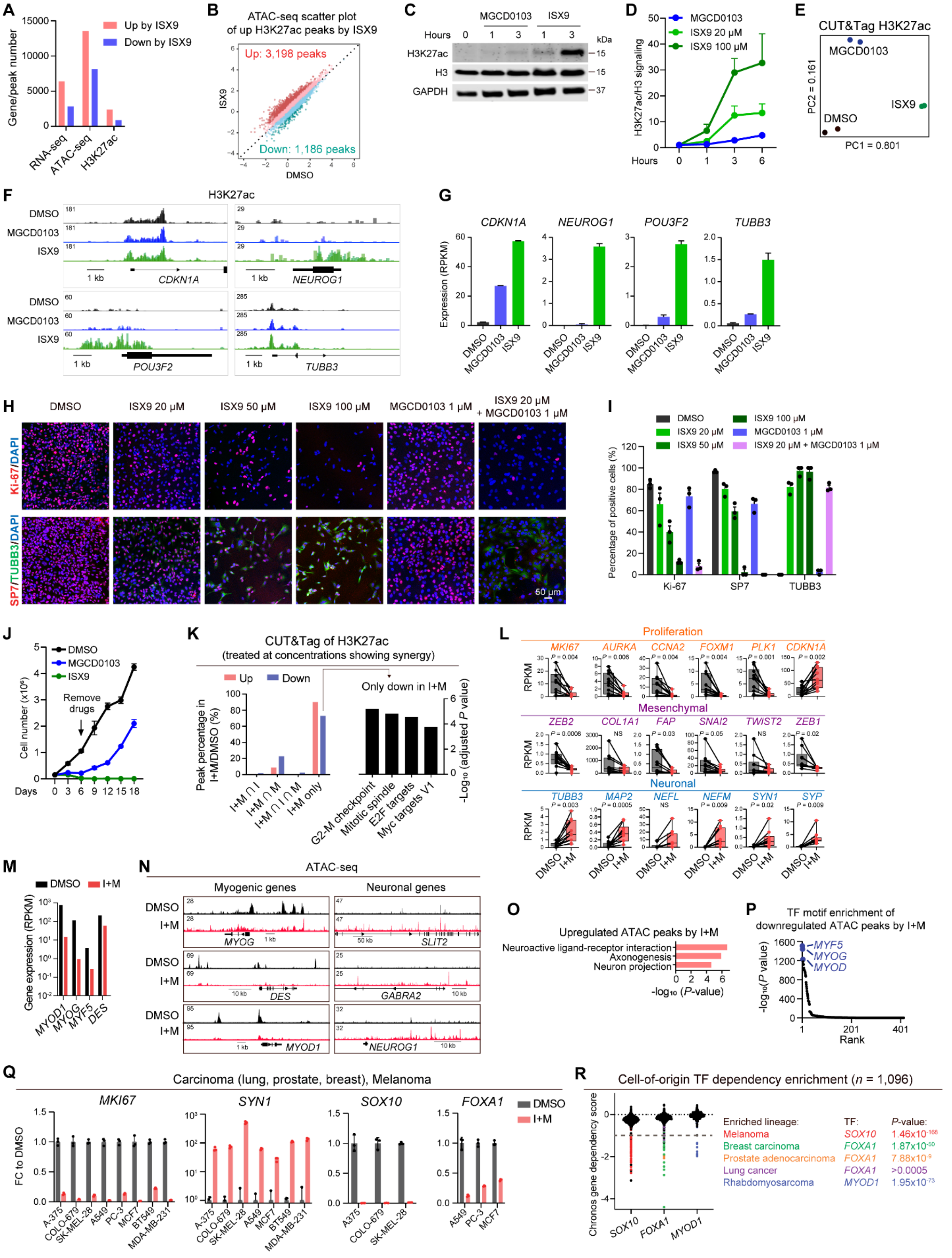

**Figure S5. Derepressing nPDC synergizes with HDAC inhibition to reprogram cancer cells, related to Figure 5.**

(A) Number of upregulated and downregulated DEGs in RNA-seq, peaks in ATAC-seq and H3K27ac peaks in CUT&Tag by ISX9 in PDXC-OS052.

(B) Scatter plot of ATAC-seq levels ($log_2$RPKM) at the H3K27ac peaks increased by ISX9. $n$ = 2.

(C) Western blot analysis of H3K27ac activation at 1 and 3 hours. MGCD0103: 1 µM. ISX9:100 µM.

(D) Quantification of the time course analysis of H3K27ac. $n$ = 3.

(E) PCA of CUT&Tag H3K27ac under 1 µM MGCD0103 and 100 µM ISX9 treatment for 3 hours.

(F) Genomic regions of the representative genes with increased H3K27ac mark. $n$ = 2 tracks overlaid.

(G) Gene expression of representative upregulated genes. $n$ = 2.

(H) Immunofluorescence staining of Ki-67 and co-staining of SP7+TUBB3 under indicated conditions. Scale bar, 50 µm.

(I) Quantification of positive cells in Figure S5H. $n$ = 3.

(J) Cell growth of PDXC-OS052 under 1 µM MGCD0103 and 100 µM ISX9 treatment ($n$ = 3). Drugs were removed on day 6.

(K) Percentage of intersection (∩) between differential H3K27ac peaks under indicated treatments compared to DMSO (left) and enrichment analysis of 3,633 exclusively downregulated H3K27ac peaks by I+M (right).

(L) RNA-seq expression of proliferation, mesenchymal, and neuronal genes under I+M and DMSO treatment in 10 sarcoma cell lines.

(M) RNA-seq of myogenic gene expression under I+M treatment for 6 days.

(N) Genomic tracks of normalized ATAC-seq densities at myogenic and neuronal genes in Rh30. $n$ = 2.

(O) Top enriched pathways among genes closest to accessible chromatin regions changed by I+M in Rh30.

(P) Enrichment of TF motif among decreased ATAC peaks by I+M.

(Q) qPCR analysis of representative proliferation, cell-of-origin and neuronal genes in melanoma (A-375, SK-MEL-28, COLO-679), lung (A549), prostate (PC-3), and breast (MDA-MB-231, BT549, MCF7) cancer cell lines under DMSO and I+M treatment. $n$ = 3.

(R) Cell-of-origin TF dependency plot showing Chronos score and significance in 1,096 cell lines from DepMap. Score of -1 indicates the median of all common essential genes. $P$-values were extracted from the DepMap database.

Data are presented as mean ± SEM (D, G, I, J, Q) or box plots divided by median and Tukey-style whiskers (L). Statistical significance was determined using Benjamini-Hochberg method (K, O) or binomial test (P).

#### Supplemental Figure 6

A

| Disease type | ID | Alternate ID | Primary Site | Resection Site | Age | Sex | Tumor type | Treatment | Mutation features | *TP53* status |
|---|---|---|---|---|---|---|---|---|---|---|
| Osteosarcoma | OS052 | PSS008 | fibula | fibula | 17 | Female | Resection | Post- | *PTEN* loss | Loss; structural variant |
| | OS152 | PSS085 | femur | scapula | 11 | Female | Metastasis | Post- | *MYC* amplification | Loss; missense variant |
| | OS525 | PSS050 | femur | lung | 11 | Female | Metastasis | Post- | *AKT1* amplification | Loss; structural variant |
| | OS526 | PSS112 | tibia | tibia | 13 | Male | Diagnostic | Pre- | *CCNE1* amplification | Loss; structural variant |
| Ewing sarcoma | EWS290 | PSS090 | sacrum | sacrum | 19 | Male | Diagnostic | Pre- | *EWS-ERG* fusion | - |
| | EWS536 | PSS159 | pelvis | pelvis | 29 | Male | Diagnostic | Pre- | *EWS-FLI* fusion | - |

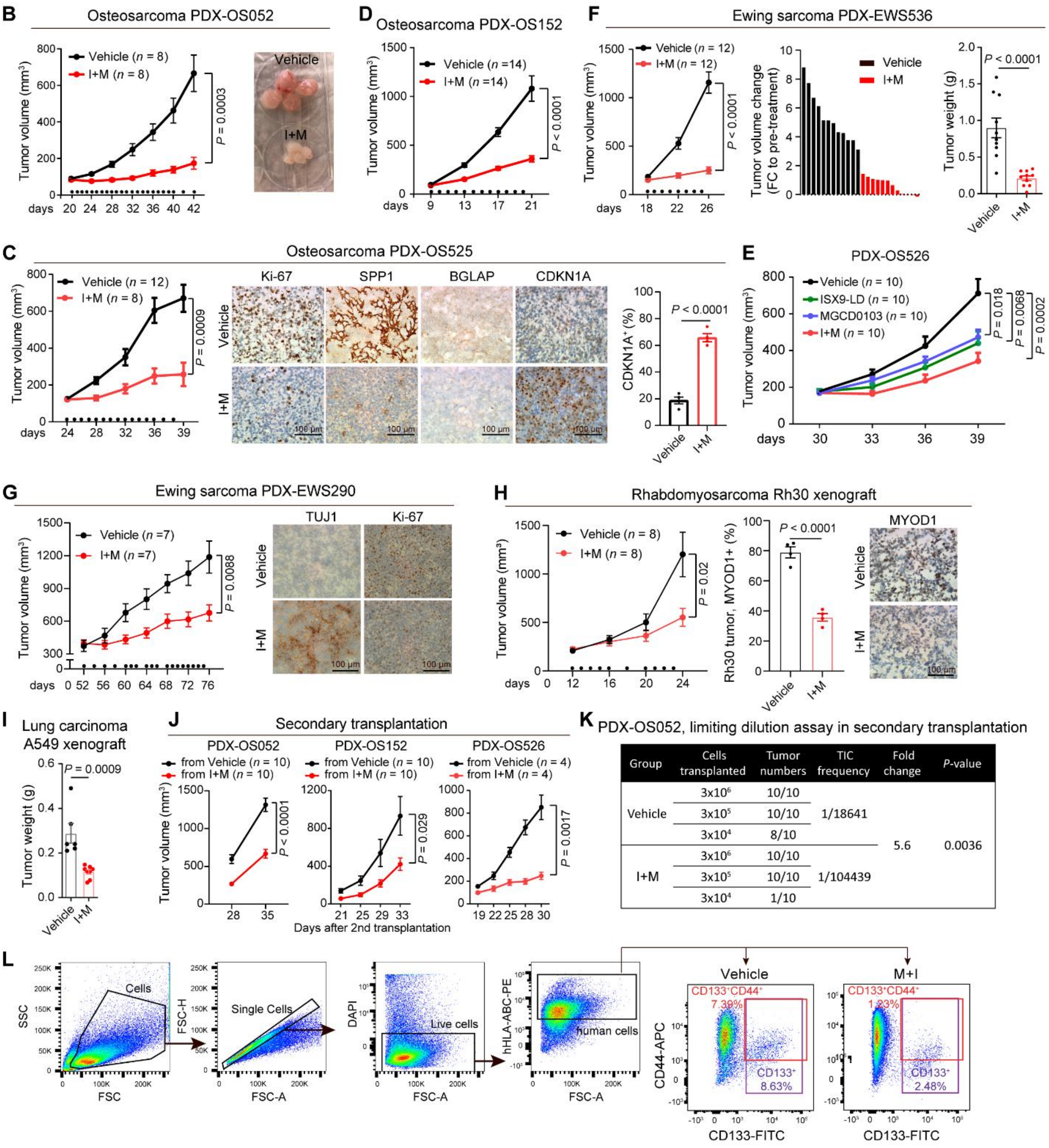


| Group | Cells transplanted | Tumor numbers | TIC frequency | Fold change | *P*-value |
|---|---|---|---|---|---|
| Vehicle | $3x10^6$ | 10/10 | | | |
| | $3x10^5$ | 10/10 | 1/18641 | | |
| | $3x10^4$ | 8/10 | | 5.6 | 0.0036 |
| I+M | $3x10^6$ | 10/10 | | | |
| | $3x10^5$ | 10/10 | 1/104439 | | |
| | $3x10^4$ | 1/10 | | | |

**Figure S6. Therapeutic cancer cell reprogramming *in vivo*, related to Figure 6.**

(A) Information on osteosarcoma and Ewing sarcoma PDX models, including ID, alternate ID, primary site, resection site, age, sex, tumor type, treatment status, mutation features, and *TP53* status.

(B) PDX-OS052. Left: tumor growth under vehicle and I+M treatments. Treatment started on day 20 post-transplantation. Middle: the image of representative tumors on a 6-well plate. Right: survival curve (determined by total tumor volume of 400 $mm^3$). All mice were euthanized on day 42 for analysis. $n = 8$ tumors. $n = 4$ mice.

(C) PDX-OS525. Tumor growth curve (left, $n \geq 8$) and IHC images of Ki-67, SPP1, BGLAP, and CDKN1A (middle, scale bar, 100 µm) and CDKN1A positive quantification (right, $n = 4$) under vehicle and I+M administration.

(D) Tumor growth curve of PDX-OS152. $n = 14$.

(E) Tumor growth curve of PDX-OS526 under daily treatment of MGCD0103, ISX9-LD, and their combination I+M. $n = 10$.

(F) Ewing sarcoma PDX-EWS536. Tumor growth curve (left), waterfall plot of individual tumor volume change (middle), and tumor weight (right) under vehicle and I+M administration. $n = 12$.

(G) Ewing sarcoma PDX-EWS290. Tumor growth curve (left) and representative IHC images of TUBB3 and Ki-67 (right) under vehicle and I+M administration. $n = 7$. scale bar, 100 µm.

(H) Rhabdomyosarcoma Rh30 orthotopic xenograft. Tumor growth curve (left, $n = 8$), quantification of MYOD1 positive cells (middle, $n = 4$), and representative IHC images of MYOD1 (right, scale bar, 100 µm) under vehicle and I+M administration.

(I) Tumor weight of lung cancer A549 xenograft under vehicle and I+M administration on day 40. $n = 6$ for vehicle and $n = 8$ for I+M. One vehicle-treated mouse was euthanized on day 37 due to ulceration.

(J) Tumor growth curve in secondary transplantation with PDX-OS052, PDX-OS152, and PDX-OS526 tumors treated with vehicle and I+M in the primary transplantation.

(K) Limiting dilution analysis of TIC in secondary transplantation by vehicle and I+M treatment in PDX-OS052.

(L) Gating strategy and the representative flow plots of TIC percentage in PDX-OS052 tumors after treatment with vehicle and I+M in Figure 7H.

B-G: tumors were transplanted on day 0, and dots on the *x*-axis represent the time points of drug administration. Data are presented as mean ± SEM. Statistics: two-tailed unpaired t-tests (B-D, F-J) or one-way ANOVA with Tukey's multiple comparison analysis (E).

#### Supplemental Figure 7

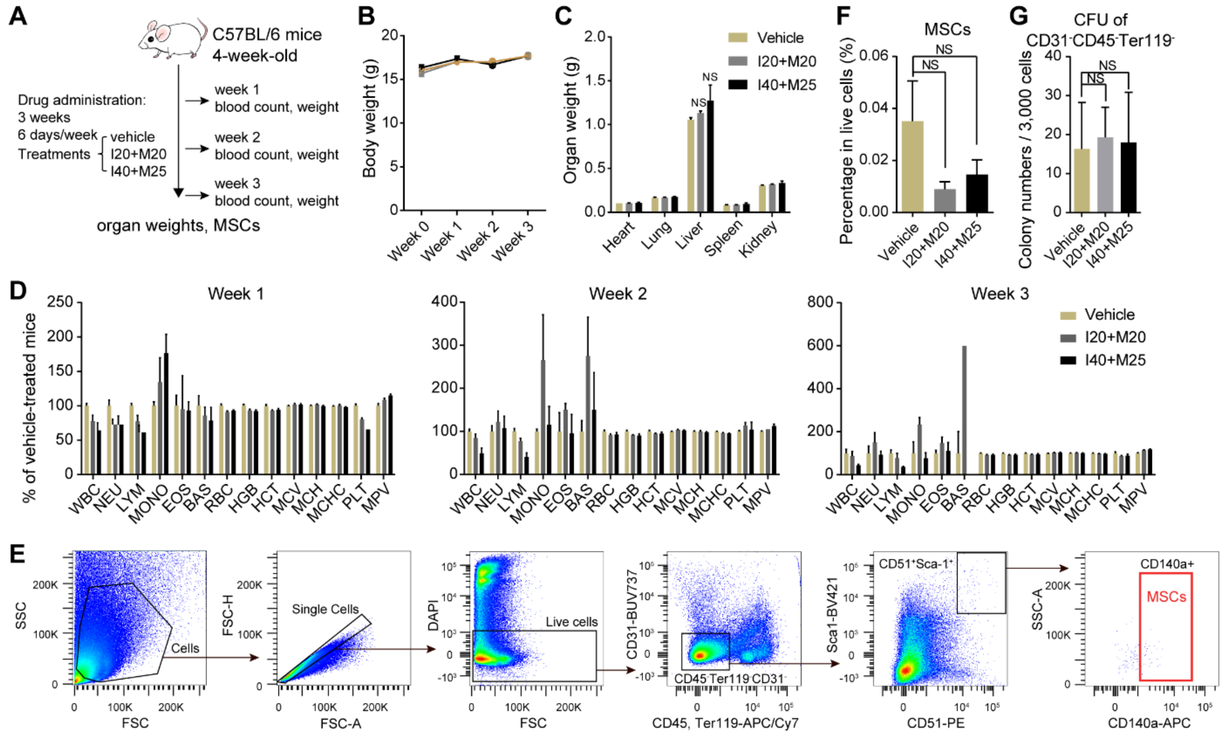


**Figure S7. Toxic studies *in vivo*, related to Figure 6.**

(A) Schematic of toxicity study *in vivo*. C57BL/6 mice were treated with I+M for 3 weeks (6 days per week) at I20+M20 (20 mg/kg ISX9 and 20 mg/kg MGCD0103) or I40+M25 (40 mg/kg ISX9 and 25 mg/kg MGCD0103).

(B) Mouse body weights. *n* = 3.

(C) Weights of vital organs at week 3. *n* = 3.

(D) Complete blood count of peripheral blood collected every week. *n* = 3.

(E) Gating strategy for sorting MSCs.

(F) Percentage of MSCs (CD31$^-$CD45$^-$Ter119$^-$Sca1$^+$CD51$^+$CD140a$^+$) in live cells isolated from bone marrow. *n* = 3.

(G) Colony formation unit assay of CD31$^-$CD45$^-$Ter119$^-$ cells counted on day 26. *n* = 3.

Data are presented as mean ± SEM. Statistics: one-way ANOVA with Tukey's multiple comparison analysis (C, F, G).
